# Environmental and transcriptomic determinants of drought response in critically endangered Siamese rosewood

**DOI:** 10.64898/2025.12.14.692107

**Authors:** Tin Hang Hung, Kalyani Lenton, Phourin Chhang, Voradol Chamchumroon, Bansa Thammavong, Riina Jalonen, Ida Theilade, John J. MacKay

**Affiliations:** Department of Biology, University of Oxford, Oxford OX1 3EL, United Kingdom; Museum of Climate Change, The Chinese University of Hong Kong, Hong Kong; Institute of Forest and Wildlife Research and Development, Phnom Penh, Cambodia; The Forest Herbarium, Department of National Park, Wildlife and Plant Conservation, Ministry of Natural Resources and Environment, Bangkok 10900, Thailand; National Agriculture and Forestry Research Institute, Forestry Research Center, Vientiane, Laos; Bioversity International, 43400 UPM Serdang, Malaysia; Department of Geosciences and Natural Resource Management, University of Copenhagen, Rolighedsvej 23, 1958 Frederiksberg C, Denmark

## Abstract

Rosewoods account for up to 40% of the global illegal wildlife trade, with *Dalbergia cochinchinensis* (Siamese rosewood) being the most heavily exploited species in Southeast Asia. Its survival is further threatened by intensifying drought linked to climate change and hydrological alteration. Here we combine greenhouse drought experiments across six provenances with full-length cDNA-seq to uncover how water-relations and carbon-use strategies vary within this species. Multivariate trait analysis resolves a two-dimensional isohydry space, in which a water-flux stringency axis (*g_s_*–*E*) is largely orthogonal to a carbon-economics axis (*A–WUE_i_*). Provenances differed strikingly, where two (KKH and DN) showed a rare *E*↓*A*↑ response, achieving high *WUE_i_* and maintaining growth under drought. Contrary to expectation, precipitation of the wettest month, not the driest, predicted isohydry, indicating that wet-season conditions set a developmental and hydrological floor for later drought responses. We identified 76 drought-responsive genes and two genes associated with isohydry axes, SEOR1 and a poorly characterised Notch-like protein AT4G14746. We also detected provenance-specific isoform switches, where drought favoured a loss-of-function PRX52 isoform lacking its signal peptide in the anisohydric provenance THB, and gain-of-function isoforms of ANN3 and LTPG5 in NP. These results reveal previously hidden diversity in drought strategies, identify mechanism-related markers for screening, and provide a simple climatic lever for climate-adjusted provenancing. We reveal post-transcriptional regulation as a novel candidate substrate for local adaptation in a threatened tropical tree, directly linking ecophysiology, climate, and genomics for conservation and restoration.

## Introduction

Extreme droughts have emerged as a driver of accelerated forest mortality globally^1,2^, which threatens terrestrial biodiversity, climate forcing, and resource availability^3^. Mechanistically, tree mortality reflects interacting processes such as xylem hydraulic failure, carbon depletion from prolonged stomatal closure, and heightened susceptibility to pests and pathogens, whose prevalences are expected to increase as warming intensifies atmospheric water demand^4^. The IPCC’s Sixth Assessment Report concludes that many regions have already experienced anthropogenically influenced increases in agricultural and ecological drought, consistent with observed forest declines during recent heat-drought events^5^. Many tree species have evolved in response to spatial and temporal variability in water availability through local adaptation and phenotypic plasticity^6^, but our understanding of these responses must be improved to predict the impacts of environmental change and to safeguard biodiversity.

Southeast Asia is a major biodiversity hotspot with disproportionally high levels of endemism, which is experiencing unprecedented levels of drought threats due to climate and land use changes. It also has the highest proportion of vascular plants, reptiles, birds, and mammals classified as in the IUCN Red List^7^. Its main water body, the Mekong River Basin, has had record low flows and severe multi-year drought in the recent decade, with widespread soil-moisture deficits, delayed monsoons and saline intrusion in the delta^8^. Drought impacts are particularly amplified by strong climate variability, such as the 2015–2016 El Niño that brings exceptional heat and rainfall deficits, with recurrent moisture shortfalls following the subsequent years^9^. The extreme drought affected area has doubled since 1950s and both the intensity and frequency of drought risks will continue to increase in all scenarios except for the lowest emission pathway^10^. Simultaneously, these climatic stresses now interact with one of the world’s fastest hydropower build-outs^11^, with more than 1,000 dams already in place with further expansion planned^12^. The large storage of water behind dams during the wet season exacerbates the downstream impacts on water availability^8^. Together these climate and infrastructure drivers are reshaping hydro-ecological regimes across Southeast Asia and underlining the need to understand adaptive drought responses for the region’s forests.

*Dalbergia cochinchinensis* Pierre (Siamese rosewood) produces extremely valuable rosewood timber in the Mekong Region, and is endemic to Cambodia, Laos, Thailand, and Vietnam. Rosewoods (*Dalbergia* spp.) amount to 30–40% of worldwide illegal wildlife trade, which is valued in total at USD 7–23 billion annually, making it the target of the world’s largest wildlife crime^13^. Siamese rosewood was assessed as ‘Critically Endangered’ in 2022 due to an estimated ≥90% population decline and population fragmentation caused by illegal logging and habitat loss^14^. The species is on CITES Appendix II since 2013, but cross-border trafficking has continued to be a significant threat, especially in Cambodia, Laos and Vietnam^14^. In addition to overexploitation, habitat conversion, fire and overgrazing, vulnerability analyses further indicate that a large share of the species range is exposed to medium to very high combined pressures, and a measurable fraction to climate-change risk by mid-century^15^. Conservation actions initiated in the early 2000s included *in situ* and *ex situ* conservation stands and seed production areas, but they were limited in scale, usually with < 50 seed-producing trees per country^16,17^. Renewed efforts to conserve the remnant populations and their genetic diversity since the 2010s have involved the collection of genetic materials, development of tree nurseries, and generation of value chains to incentivise local livelihoods^18,19^.

With increasing drought risks in Mekong Region, our ability to safeguard the survival and conservation tree species such as Siamese rosewood requires a better understanding of the variation in drought resistance and growth performance.

A useful lens to help fill this gap is the isohydry-anisohydry continuum. Isohydric plants stabilise leaf water potential (Ψ_leaf_) by closing stomata early as soils dry, while anisohydric plants allow Ψ_leaf_ to decline to maintain carbon assimilation. We previously reported that Siamese rosewood had an anisohydric response to short-term drought, which maximised assimilation at the cost of water loss, whereas a isohydric response was found in the sympatric species *D. oliveri*^20^. While this anisohydric response can sustain photosynthesis during drought, it increases hydraulic risk and may lead to mortality under prolonged water deficits^21^.

Identifying the molecular mechanisms and key genes in drought will provide additional insights for the physiological responses. Recent population genetic studies have shown that Siamese rosewood is predominantly outcrossing and thus retains high levels of genetic diversity across much of its range despite severe demographic declines^22^. Central populations in Cambodia and eastern Thailand harbour the highest allelic richness, whereas peripheral and heavily exploited stands in northeastern Thailand, Laos, and Vietnam tend to exhibit reduced diversity and higher relatedness, consistent with historical bottlenecks, habitat fragmentation, and local exploitation^23^. In addition, our previous genomic scan has identified substantial genetic differentiation in *D. cochinchinensis* driven by temperature- and precipitation-related environmental factors^24^. Local adaptation is also found to be the strongest in the periphery of the species range^24^. Thus, it is very plausible that populations will display differentiated response to drought. However, it has also been shown that environment has a stronger effect on gene expression than standing genetic variation^25^. The gene expression or transcriptomic response can be considered an intermediate phenotype that is informative of high-level physiological traits^26^. It is well-established that plants remodel their transcriptome under drought stress, which regulates stress sensors, signalling pathways, and synthesis of hormones and enzymes^27,28^. However, fundamental research in forest trees is still underrepresented compared to other plant crops^29^, and systematic studies at population level are scarce^30^.

The endangered status of and the drought threats facing Siamese rosewood together highlight the need to improve our understanding of its drought tolerance variability and adaptation across the remnant populations. The overarching aim of this study is to identify the environmental, physiological, and transcriptomic determinants of the species-wide heterogeneity in drought response, by building on the recent capacity in genomic research in Siamese rosewood, including its high-quality reference genome and range-wide genomic scan^24,31^. Seedling recruitment is a critical bottleneck in Siamese rosewood^32^ and early phenotyping can still provide valuable insights into drought adaptation. First, we characterise the drought response in seedlings from six provenances by comparing various physiological traits in a greenhouse drought experiment. Second, we identify genes that are differentially expressed among different provenances under drought stress. Third, we analyse if different provenances have differential transcript usage in response to drought. This study will ultimately ensure drought resilience in germplasm of Siamese rosewood by integrating recent developments in conservation and cutting-edge genomic technologies.

## Methods

### Plant materials

Dried seeds of *Dalbergia cochinchinensis* were provided by the Institute of Forest and Wildlife Research and Development, Cambodia; the National Agriculture and Forestry Research Institute, Laos; and the Department of National Park, Wildlife and Plant Conservation, Thailand in 2020 from six local seed sources across the species range (**Figure 1a** and **Supplementary Table 1**). The seeds were scarified by placing them in 70°C distilled water and left to cool to room temperature overnight. They were germinated on 1% agar in a controlled greenhouse at 30°C and photoperiod 12L/12D. Germinants were first transferred to 0.125L pots in a soil-perlite 3:1 (v:v) mixture and grown for three months. Healthy seedlings were then transferred to 0.81 L pots with the same substrate and grown for further two months. Throughout this period, plants were watered regularly to maintain at substrate capacity and fertilized once a week using NLPLK 20:20:20 fertilizer (Chempak, Suffolk, United Kingdom).

**Figure 1.**
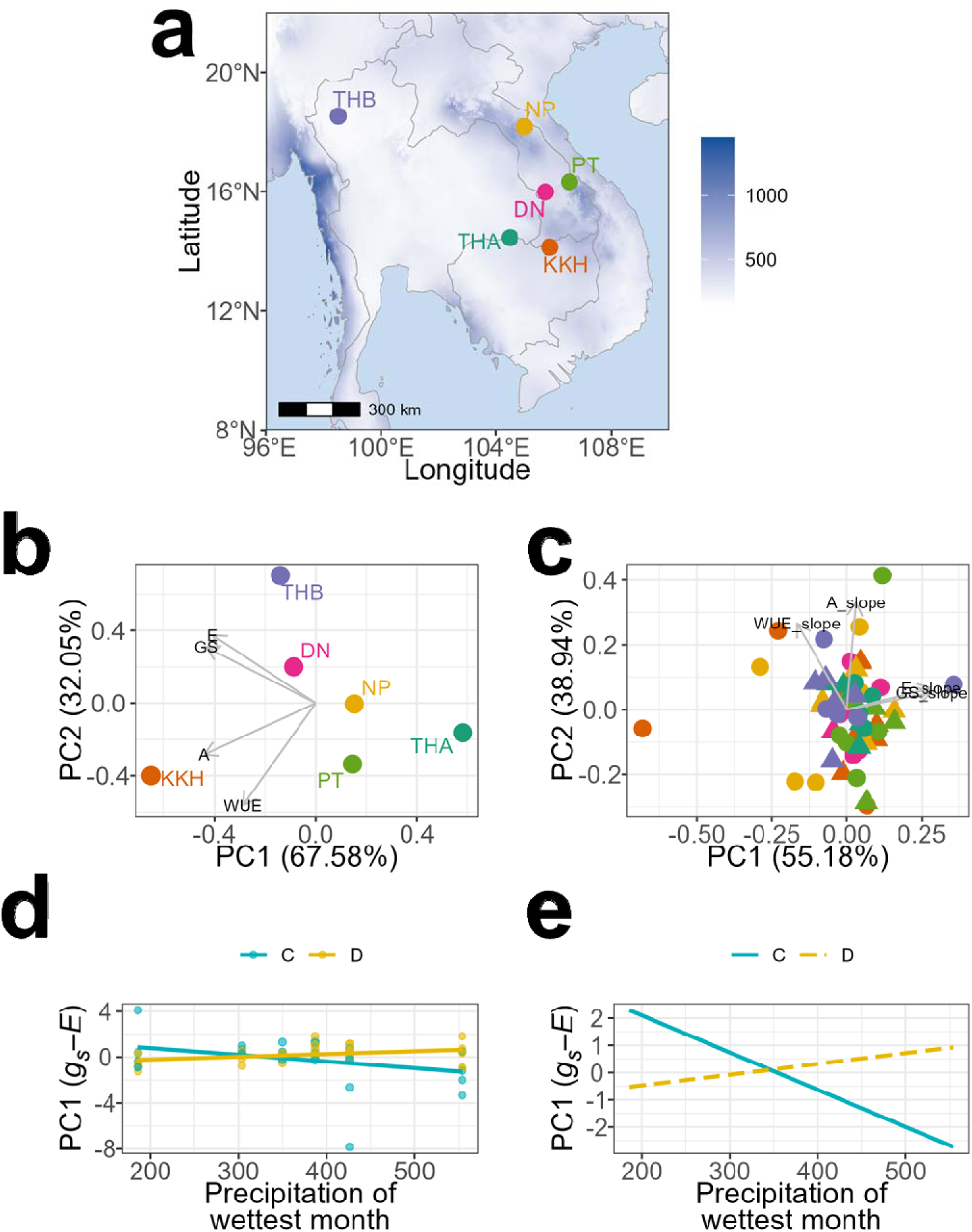
**(a)** Six local seed provenances of *Dalbergia cochinchinensis* in this study. The map is colour coded with the precipitation of wettest month (mm). The colour scheme of the provenances is consistent in all figures throughout this manuscript. **(b)** Provenance-level PCA on the coefficients of effect of Treatment × Provenance on physiological traits that show significant interactions. **(c)** Individual-level PCA on physiological traits that show significant interactions. **(d)** The interaction effect of precipitation of wettest month and treatment on the isohydry score (which is a composite score derived from the first principal component (PC1) of the individual-level PCA in Figure 1c, see **Methods** for details). **(e)** Corresponding interaction plot, which shows model-predicted relationships and the fitted values between precipitation of wettest month and isohydry score.

**Figure 2.**
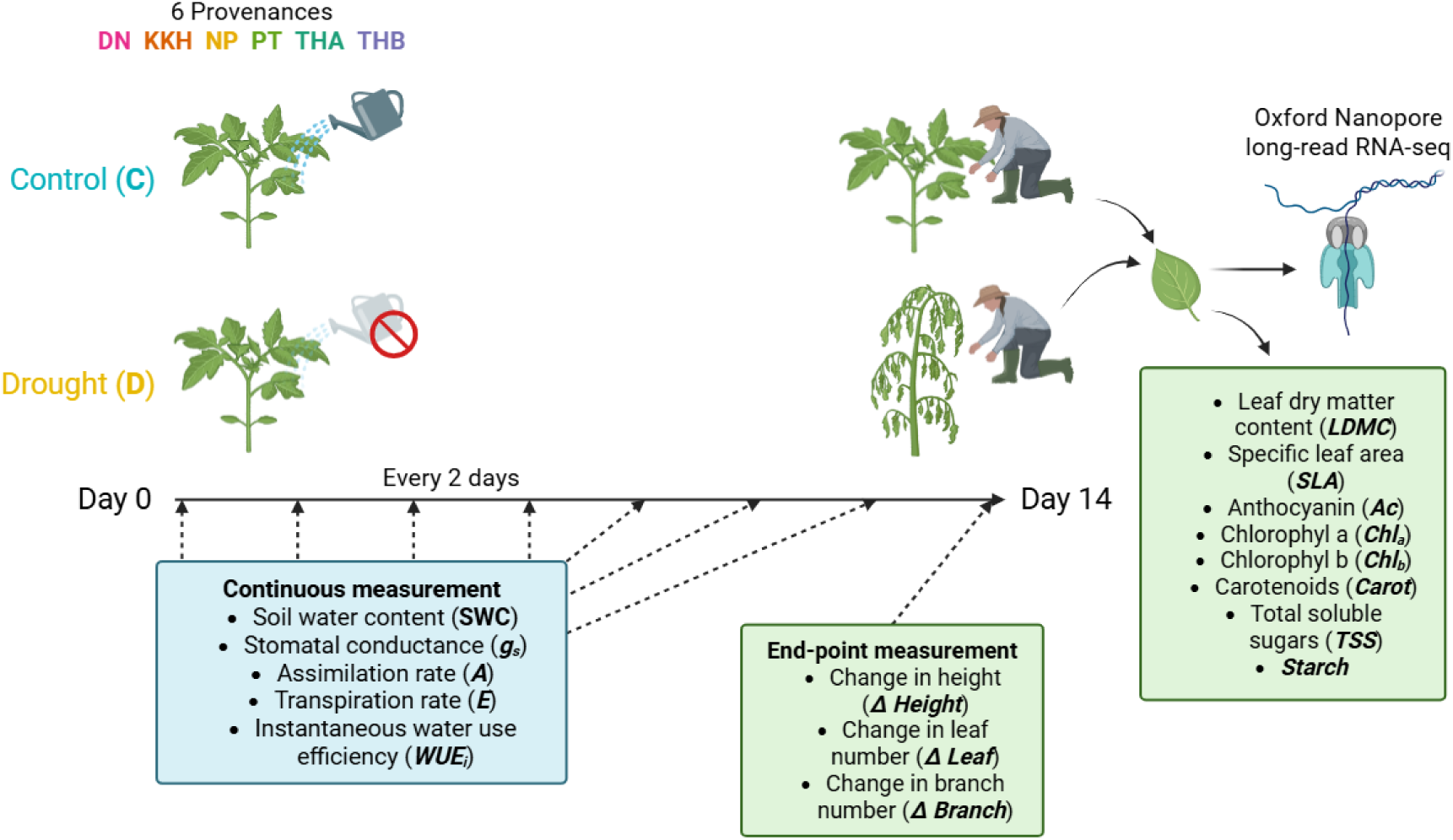
Experimental design of the greenhouse experiment. We studied 10 plants per provenance and randomly assigned half of the plants to either well-watered control (C) or water-withholding treatment (D).

### Experimental design

We used a split-plot design with 10 plants per provenance randomly distributed into 10 trays, making up to 60 plants in total. We randomly assigned half of the trays to either well-watered control (C) or water-withholding treatment (D). The controls were watered every other day to maintain substrate capacity at 40–55%, whereas the droughted were not watered at all after the experiment started. We also randomly split the trays into two blocks with the start of experiment staggered by one day.

There were two types of data collected: (1) continuous data of water relation and photosynthetic measurements, and (2) end-point data of anatomical traits and biochemical measurements. At the end of the experiment (day 14), three leaves were sampled from each plant, snap-frozen in liquid nitrogen, and stored at –80°C.

### Water relation and photosynthetic measurements

Measurements were taken between 10 am and 2 pm (at least two hours after sunrise and two hours before sunset), when photosynthetic activity reached its plateau (i.e. point of saturation). We measured soil water content (*SWC*) by using a ML3 ThetaProbe Soil Moisture Sensor (Delta-T Devices Ltd., Cambridge, England). We measured stomatal conductance (*g_s_*), photosynthetic assimilation rate (*A*), and transpiration rate (*E*) using an infrared gas analyser LCpro T, Leaf Chamber & Soil Respiration System (SRS2000 T, ADC BioScientific Limited, England) and its broad leaf chamber, with the light intensity set to PAR 500 with equal parts of red, green, and blue lights. We calculated instantaneous water use efficiency (*WUE_i_*) using the following equation^33^:

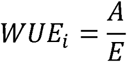

### Anatomical traits

We recorded changes in height (Δ *Height*), leaf number (Δ *Leaf*), and branch number (Δ *Branch*) before and after the experiment. We measured leaf area using the equation for the area of an ellipse (*S* = *ab*π). We measured fresh (*FW*) and dry weights (*DW*) of the leaves before and after lyophilisation of two days in an Alpha 2-4LD-2 laboratory freeze-dryer (Martin Christ GmbH, Germany). We calculated the leaf dry matter content (*LDMC*) and specific leaf area (*SLA*) using the following equations^34^:

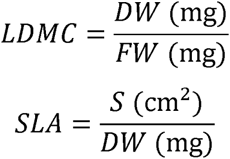

### Pigment extraction and quantification

We ground ∼25 mg of lyophilised leaves with a TissueLyzer (Retsch, Germany) at 25^−1^ s for 1 minute. We then extracted the pigments by adding an 80:20 mixture (v:v) of cold acetone and 50 mM Tris buffer pH 8.0 and incubated for 72 hours, following Sims & Gamon’s protocols^35^. After centrifugation, we transferred the supernatant to a 15ml Falcon tube and then topped the extracts up with 6–12 ml of the acetone-Tris buffer to ensure that the absorbances below were approximately in the range between 0 and 1. We used a Nanodrop One (Thermo Fisher Scientific, Waltham, Massachusetts, USA) to measure the absorbances of the extracts containing the leaves at 470, 537, 647, and 663 nm. We determined the concentrations of anthocyanin (*Ac*), chlorophyll a (*Chl_a_*) and b (*Chl_b_*), and carotenoids (*Carot*) with the following equations:

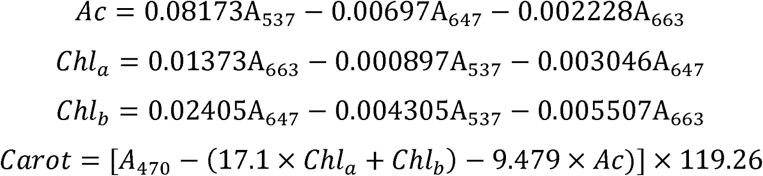

### Sugar extraction and quantification

We heated ∼25 mg of lyophilised leaves in 80% ethanol for one hour at 80°C. After centrifugation, we determined glucose concentration from the supernatant using Osaki’s anthrone method^36^. We mixed the supernatant with anthrone and sulfuric acid and heated it at 100°C for 10 minutes sharply. We cooled the mixture on ice and measured their absorbances at 625 nm using a Genesys 150 UV-Visible spectrophotometer (Thermo Fisher Scientific, Waltham, Massachusetts, USA). We calculated the concentration of total soluble sugars (*TSS*) in the samples according to a D-glucose standard curve.

We washed the pellets in 80% ethanol to remove any glucose and then heated them at 100°C for 10 minutes with 30% perchloric acid to convert all starch to glucose. We again measured absorbances of the extract at 625 nm again and used a D-glucose standard curve to determine the concentration of starch (*Starch*).

### Statistical analyses on traits

All statistical analyses were conducted in R 4.5.1. List of all traits included in this analysis is presented as **Supplementary Table 2.**

We applied square-root transformation to *g_s_*, *A*, *E*, and *WUE_i_* and logarithmic transformation to *Ac*, *Chl_a_*, *Chl_b_*, and *Carot* to correct for normality. We assessed normality based on the distribution of the residuals using a QQ plot. For continuous data, we conducted two-way ANOVAs to examine the effects of *SWC*, provenance, and their interaction term on *g_s_*, *A*, *E*, and *WUE_i_*. For end-point data, we conducted two-way ANOVAs to examine the effects of treatment, provenance, and their interaction term anatomical and biomass traits (Δ *Height*, Δ *Leaf*, Δ *Branch*, *LDMA*, *SLA*), pigments (*Ac*, *Chl_a_*, *Chl_b_*, and *Carot*), and sugars (*TSS* and *Starch*). We initially included the blocks as a random variable in the ANOVAs, but no difference was found thus the random variable was excluded from the models.

### Quantification and climatic associations of isohydry

We quantified the provenance-level isohydry based on the principal component analysis (PCA) of the coefficients of interaction effect of treatment × provenance on physiological traits that show significant interactions. We then quantified individual-level isohydry also based on physiological traits that show significant interactions, with *g_s_*, *A*, *E*, and *WUE_i_* divided by *SWC* to obtain a single-value slope for each individual. Visual inspection confirmed that the loading of treatment aligned with the direction of second principal component (PC2), thus the first principal component (PC1) was used as a composite proxy for isohydry (denoted as *Isohydry PC1* in this manuscript).

Provenance-level PCR is useful in quick visualisation and quantification of isohydry at whole-provenance level, which is more spatially explicit and useful in conservation context. However, individual-level PCR reveals a more substantial variation in isohydry among individual trees that are not attributable to provenances, and thus is more suitable for subsequent analyses on environmental and transcriptomic associations.

We tested the effect of 19 climatic variables that were biologically meaningful, from the WorldClim 2 database (*bio_1–bio_19*). To correct for the multicollinearity inherent in climatic data, we used a stepwise approach to remove climatic variables that have a variance inflation factor (VIF) > 10. We then examined the effects of treatment, these climatic variables, and their interactions on both Isohydry PC1 and PC2, using multivariate analysis of variance (MANOVA).

### RNA extraction, full-length cDNA library construction, and sequencing

We isolated total RNA from leaves using the Monarch Total RNA Miniprep Kit (New England Biolabs, United Kingdom). We determined their quantity on a Qubit 4 Fluorometer (Thermo Fisher Scientific, United Kingdom), assessed their purity using a NanoDrop One Spectrophotometer (Thermo Fisher Scientific), with A260/280 and A260/230 above 1.80, and verified their integrity on a 1% bleach-agarose gel.

The 6-µl reverse transcription reaction contained 3 µl of ∼200 ng total RNA, 2 µl of 10 µM Reverse Transcription Primer (5’–AGCAGTGGTATCAACGCAGAGTAC(T)_30_V–3’) and 1 µl of 10 mM dNTP. The reaction was incubated at 70°C for 5 min and held on ice to allow the primer to anneal to mRNA with poly(A) tail. cDNA synthesis was performed by adding 2.5 µl of Template Switching RT Buffer, 1 µl of Template Switching RT Enzyme Mix (#M0466, New England Biolabs, United Kingdom), and 0.5 µl of 75 μM Template Switching Oligo (TSO) (5’–GCTAATCATTGCAAGCAGTGGTATCAACGCAGAGTACATrGrGrG–3’) to the 6-µl primer-annealed reaction. The reaction was incubated at 42°C for 90 min, 85°C for 5 min, and held at 4°C. Full-length cDNA (fl-cDNA) was amplified by adding 12.5 µl of Q5 Hot-Start High-Fidelity 2X Master Mix (New England Biolabs, United Kingdom), 10 µl of nuclease-free water, and 1 µl of 10 µM cDNA PCR Primer (5’– AAGCAGTGGTATCAACGCAGAGT–3’) to the 10 µl cDNA product. The thermal cycling profile was 98°C 45 s, 20× [98°C 10 s, 62°C 15 s, 72°C 3 min], 72°C 5 min, for yielding ∼1 µg fl-cDNA product. We cleaned up the fl-cDNA using 1.2× AMPure XP (Beckman Coulter, United States).

Nanopore libraries were constructed with the ligation sequencing chemistry using ∼200 fmol pooled library (∼250 ng for 2,000 bp cDNA). Nanopore libraries were then sequenced and basecalled using the super-accuracy model on a GridION system (Oxford Nanopore Technologies, United Kingdom) at the Department of Biology, University of Oxford.

Basecalled reads were trimmed for Nanopore adaptors, the primer sequences, and split for chimeras using dorado 0.6.2+14a7067. We then identified and quantified known and novel transcripts using IsoQuant 3.4.1^37^ and minimap 2.28-r1209, supplied with the reference genome and gene annotation of *D. cochinchinensis* (Dacoc 1.2). We also extracted the transcript sequences from the transcript model generated by IsoQuant with gffread 0.12.7.

### Differential gene expression analysis

We imported the gene-level count data from IsoQuant into R 4.4.1 and performed the analysis with DESeq2 1.44.0^38^. We removed low-expression genes where there were less than 6 samples (the size of the smallest experimental unit) with normalised counts greater than or equal to 10. We conducted likelihood ratio tests to compare a full model which accounts for provenance, ∼ `Treatment + Provenance + PC1 + PC2 + Treatment:PC1 + Treatment:PC2`, with a reduced model in which the effect of interest is removed. First, we tested the main effect of drought treatment on gene expression (*the drought effect*) with a reduced model of ∼ `Provenance + PC1 + PC2 + Treatment:PC1 + Treatment:PC2`. Second, we tested the differential effect of treatment on gene expression in isohydry (*the isohydry effect*), as represented by the two principal axes (PC1 and PC2) using two LRTs with reduced models of ∼ `Treatment + Provenance + PC1 + PC2 + Treatment:PC2` and ∼ `Treatment + Provenance + PC1 + PC2 + Treatment:PC1` for PC1 and PC2 respectively. For both tests, we applied independent hypothesis weighting, which could increase detection power in genome-scale multiple testing^39^, with an FDR threshold of 0.05 to discover significantly differentially expressed genes (DEG).

We conducted gene set enrichment analyses (GSEA) on the effect size, which is the χ^2^-statistics in LRT, to search for Gene Ontology (GO) terms and Kyoto Encyclopedia of Genes and Genomes (KEGG) pathways that are significantly enriched using clusterProfiler 4.12.0^40^ and fgsea 1.30.0^41^.

### Differential transcript usage, annotations, and functional consequences

We analysed differential transcript usage with the pipeline IsoformSwitchAnalyzeR 2.8.0^42^, which incorporated DEXSeq 1.55.1^43^. First, we assessed isoform switching between control and drought-treated individuals, and set provenances as the batch effect (*the overall drought effect*). Second, we assessed isoform switching between control and drought-treated individuals for each provenance (*the provenance effect*). We determined significant differential transcript usage as those with a difference in isoform usage (dIF) > 0.01 and an isoform switch *Q*-value < 0.05. Individual-level isohydry could be not used as a factor, like in the case of differential gene expression analysis, because continuous variables are not compatible with existing differential transcript usage analysis pipelines.

We annotated isoforms exhibiting significant differential transcript usage using a suite of complementary tools. Coding potential was assessed with CPC2 1.0.1^44^. Conserved protein domains were identified using pfam_scan.pl on Pfam database version 38.0^45^. Signal peptides were predicted with SignalP 6.0^46^. Subcellular localisation was inferred using DeepLoc 2.0^47^ in Accurate mode. Transmembrane helices were predicted with DeepTMHMM 1.0.44^48^. Intrinsically disordered protein regions were identified using AIUPred v2.1.2^49^ (aka IUPred 3). These annotations were integrated within the IsoformSwitchAnalyzeR framework to evaluate the functional consequences of isoform switching events, including potential changes in coding potential, protein domain architecture, secretion signals, membrane localisation, and structural disorder.

## Results

### Variable responses in water relation and photosynthetic traits

All four water relation and photosynthetic traits varied significantly in response to the changes in soil water content (SWC) caused by the drought treatment. We also observed provenances effects and provenance x SWC interactions for several of the same traits. Stomatal conductance (*g_s_*) decreased with declining SWC by a coefficient of –8.063e-05 (F_1,408_ = 4.11, *p* = 0.043), with significant variation among provenances (F_5,408_ = 2.93, *p* = 0.013) and a significant SWC × provenance interaction (F_5,408_ = 2.74, *p* = 0.019). Positive slopes (higher *g_s_* with higher SWC) were observed in THA (1.20e-03), NP (1.03e-03), and PT (1.01e-03), while negative slopes (lower *g_s_* with higher SWC) were found in KKH (–5.52e-05), DN (–8.06e-05), and THB (–7.73e-04) (**Figure 3a–b**). Photosynthetic assimilation rate (*A*) also showed significant effects of SWC (F_1,408_ = 5.39, *p* = 0.021) and provenance (F_5,408_ = 4.47, *p* < 0.001), as well as a strong SWC × provenance interaction (F_5,408_ = 4.97, *p* < 0.001). THA (0.0037) and NP (0.00051) had positive slopes, while PT (–0.0014), THB (–0.0018), DN (–0.0038), and KKH (–0.010) had negative slopes (**Figure 3c–d**). Transpiration rate (*E*) was strongly influenced by SWC (F_1,408_ = 14.52, *p* < 0.001) and provenance (F_5,408_ = 4.14, *p* = 0.0011), while the SWC × provenance interaction was nearly significant (F_5,408_ = 2.06, p = 0.069). Five of the provenances produced positive slopes PT (0.0060), THA (0.0059), NP (0.0051), KKH (0.0020), and DN (0.0014) had and a negative slope was only observed in THB (–0.0014) (**Figure 3e–f**). Instantaneous water-use efficiency (*WUE_i_*) increased significantly under lower SWC (F_1,408_ = 19.82, p < 0.001), with a significant SWC × provenance interaction (F_5,408_ = 4.23, p < 0.001) but no main effect of provenance. Accordingly, most of the provenances had negative slopes, while THA (–0.0012), NP (–0.0029), PT (–0.0057), DN (–0.0065), and KKH (–0.015), while a positive slope was found for THB (0.00057) (**Figure 3g–h**).

**Figure 3.**
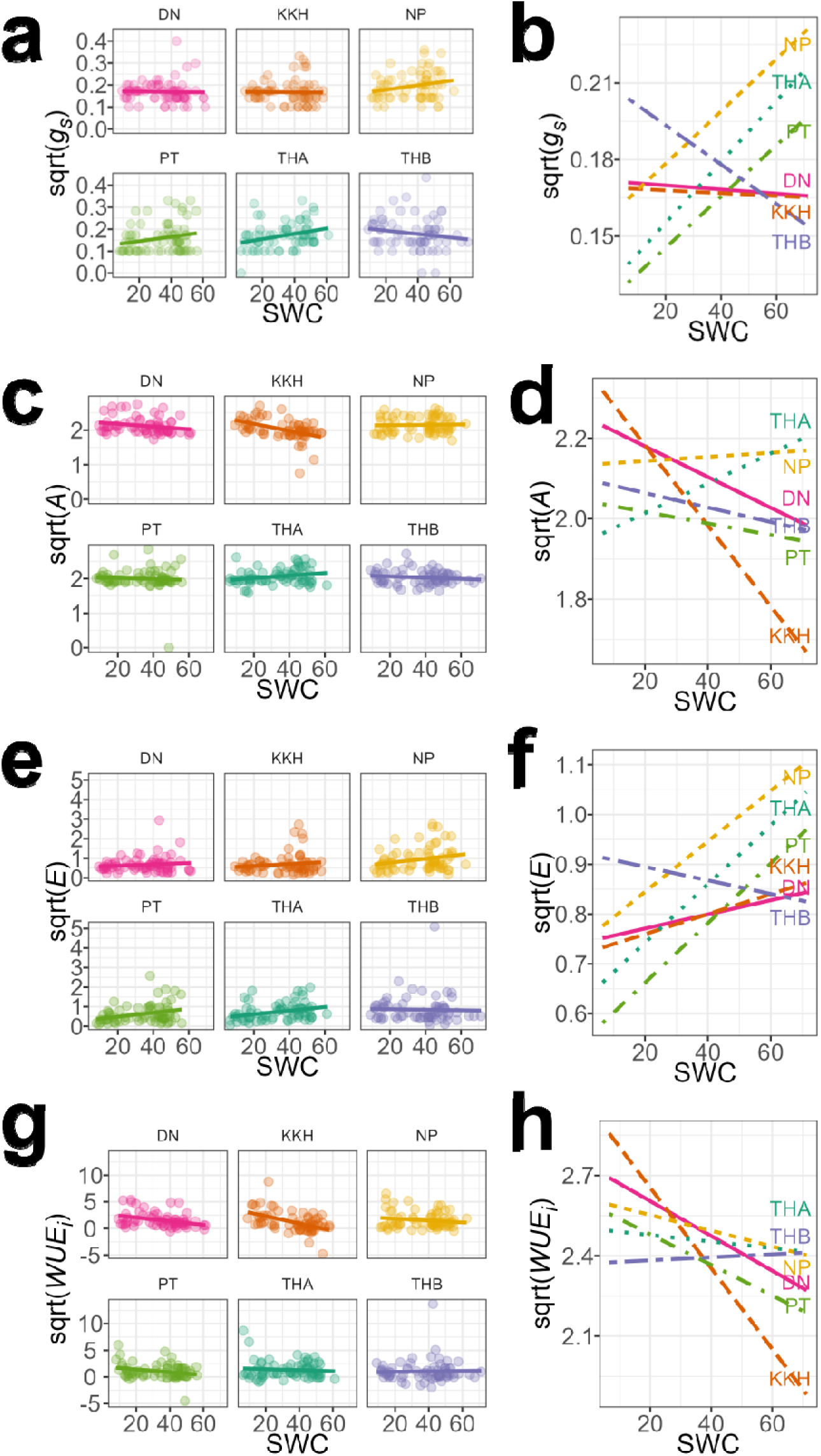
**(a)** Stomatal conductance (*g_s_*), **(c)** assimilation rate (*A*), **(e)** transpiration rate (*E*), and **(g)** water use efficiency (*WUE_i_*) along the gradient of soil water content (SWC). All traits were square-root transformed to correct for normality. **(b)**, **(d)**, **(f)**, and **(h)** Corresponding interaction plots of *g_s_*, *A*, *E*, and *WUE_i_*, which show model-predicted relationships and the fitted values between SWC and the corresponding trait.

### Anatomical and biochemical traits

Height change (Δ *Height*) showed a significant treatment × provenance interaction (F_5,48_ = 3.06, *p* = 0.0177), although treatment (F_1,48_ = 1.47, *p* = 0.23) and provenance (F_5,48_ = 1.34, *p* = 0.26) alone were not significant. Final height was higher in drought-treated individuals in KKH (+1.2 cm), DN (+0.8), and NP (+0.2), and lower in drought-treated individuals in THB (–0.8), PT (–1.2), and THA (–2.6) (**Figure 4a–b**). Leaf (Δ *Leaf*) and branch (Δ *Branch*) number changes were unaffected by treatment, provenance, or their interaction (*p* > 0.05) (**Supplementary Figure 1a–b**).

**Figure 4.**
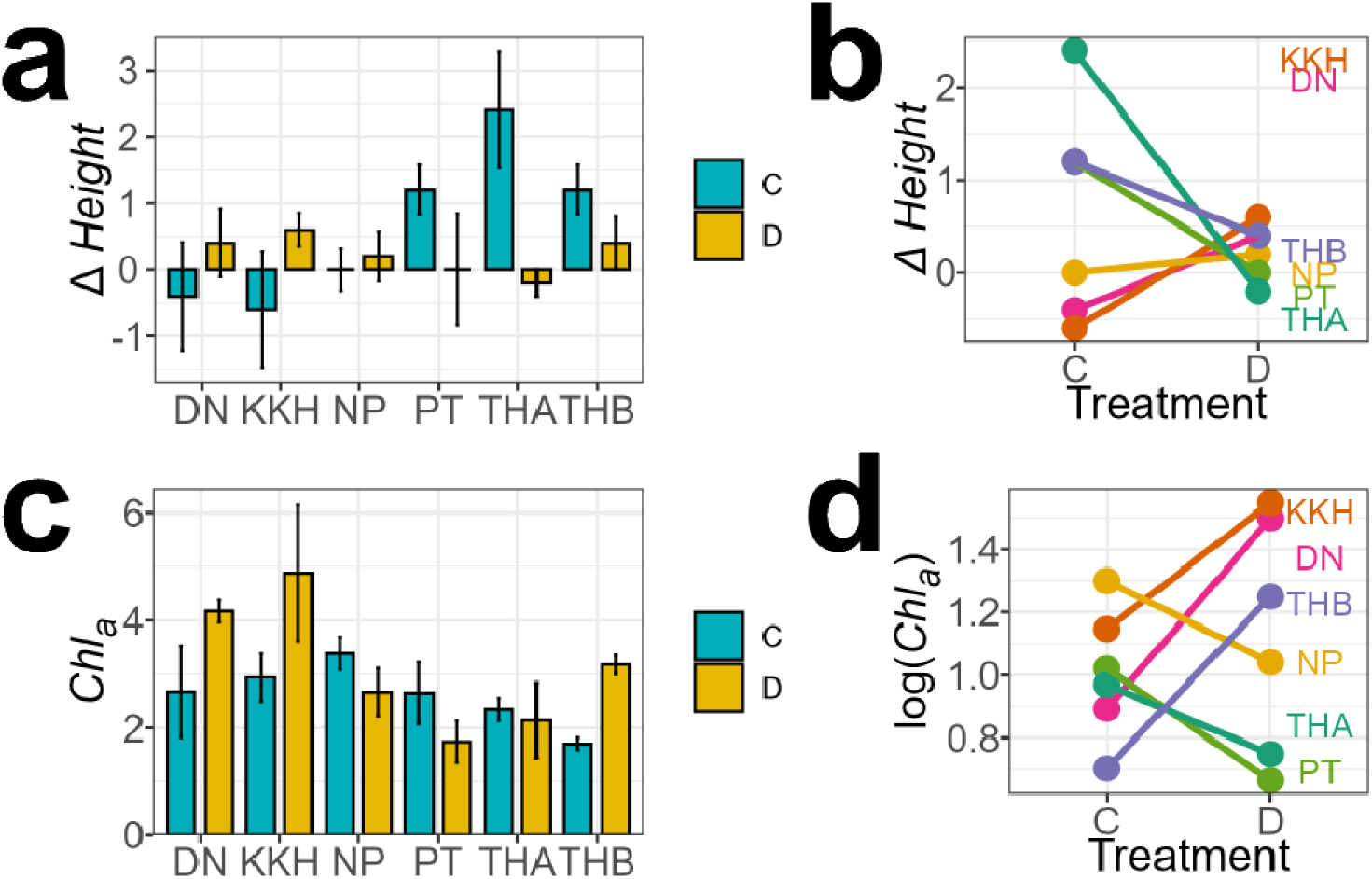
**(a)** Change in height (Δ *Height*) and **(c)** chlorophyll a content (*Chl_a_*) between control (C) and drought (D) among six provenances. **(b)** and **(d)** are the corresponding interaction plots, which show model-predicted relationships and the fitted values between treatment and the corresponding traits. *Chla* was log-transformed to correct for normality in the model in Figure 3d, but raw values were visualised in Figure 3c.

Chlorophyll a content (*Chl_a_*) showed significant effects of provenance (F_5,48_ = 2.65, *p* = 0.0342) and a significant treatment × provenance interaction (F_5,48_ = 3.14, *p* = 0.0156). *Chl_a_* was higher in drought-treated individuals in KKH (+1.94), DN (+1.50), THB (+1.48), and lower in THA (–0.20), NP (–0.73), and PT (–0.91) (**Figure 4c–d**). No significant effects were detected for anthocyanin (*Ac*), chlorophyll b (*Chl_b_*), or carotenoids (*Carot*), although the provenance effect on carotenoids was marginal (F_5,48_ = 1.99, *p* = 0.0966) (**Supplementary Figure 1c–e**).

Neither leaf dry matter content (*LDMC*) nor specific leaf area (*SLA*) was significantly affected by treatment, provenance, or their interaction, although SLA showed a marginal treatment effect (F_1,48_ = 2.99, p = 0.0904) (**Supplementary Figure 1f–g**).

Neither total soluble sugars (*TSS*) nor starch (*Starch*) concentrations differed significantly among treatments, provenances, or their interactions (*p* > 0.05) (**Supplementary Figure 1h–i**).

All ANOVA tables were summarised in **Supplementary Table 3**.

### Characterisation and environmental association of isohydry

We characterised provenance- and individual-level isohydry using the first two principal components (PC1 and PC2) that summarise the water relations and photosynthetic traits. The provenance-level two-dimensional isohydry space captured 67.58 + 32.05 = 99.63% of the variation in the coefficients of effect of SWC × Provenance on the water relations and photosynthetic traits (**Figure 1b**). PC1 co-varied largely with *A* and *WUE_i_*, while PC2 co-varied largely with *g_s_* and *E*. The co-direction of loadings suggested that *g_s_* and *E* were positively correlated, and the same between *A* and *WUE_i_*. However, the orthogonality suggested that *g_s_*–*E* and *A*–*WUE_i_* were largely independent from each other.

The individual-level two-dimensional isohydry space captured 55.18 + 38.94 = 94.12% of the variation in water relations and photosynthetic traits (**Figure 1c**). PC1 co-varied largely with *g_s_* and *E*, while PC2 co-varied largely with *A* and *WUE_i_*. Similarly, *g_s_*–*E* and *A*–*WUE_i_* were largely independent from each other.

Bioclimatic variables were largely inter-correlated and after filtering, only four variables were retained in the model (VIF < 10), namely precipitation of wettest month (*bio_13*), precipitation of driest month (*bio_14*), and precipitation seasonality (*bio_15*). Only precipitation of wettest month (*bio_13*) showed a significant treatment × isohydry interaction (F_2,51_ = 3.20, *p* = 0.049) (**Figure 1d–e**). Higher precipitation of the wettest month led to a higher isohydry score.

### Differentially expressed genes for drought response and isohydry

Full-length transcriptome sequencing yielded an average of 2.40 Gb (SD ± 0.85) for 36 individuals. The mean read length was 729.20 bp and N50 was 952.33 bp.

For the *drought effect*, we detected 76 genes that were significantly differentially expressed (**Figure 5a** and **b** and **Supplementary Table 4**). The most statistically significant, annotated genes were GASA14 (Dacoc22547), CYP77A4 (Dacoc12206), PME1 (Dacoc00037), DEG11 (Dacoc04747), CXE20 (Dacoc12653), RDUF2 (Dacoc26635), XTH9 (Dacoc27022), and THE1 (Dacoc27311). There were 21 drought-response genes that were unannotated. Five gene ontology terms were enriched, including Golgi cis cisterna (GO:0000137) (*q* = 0.0017), cell wall organization or biogenesis (GO:0071554) (*q* = 0.016), small molecule biosynthetic process (GO:0044283) (*q* = 0.016), external encapsulating structure organization (GO:0045229) (*q* = 0.019), and cell wall organization (GO:0071555) (*q* = 0.025) (**Supplementary Table 5**).

**Figure 5.**
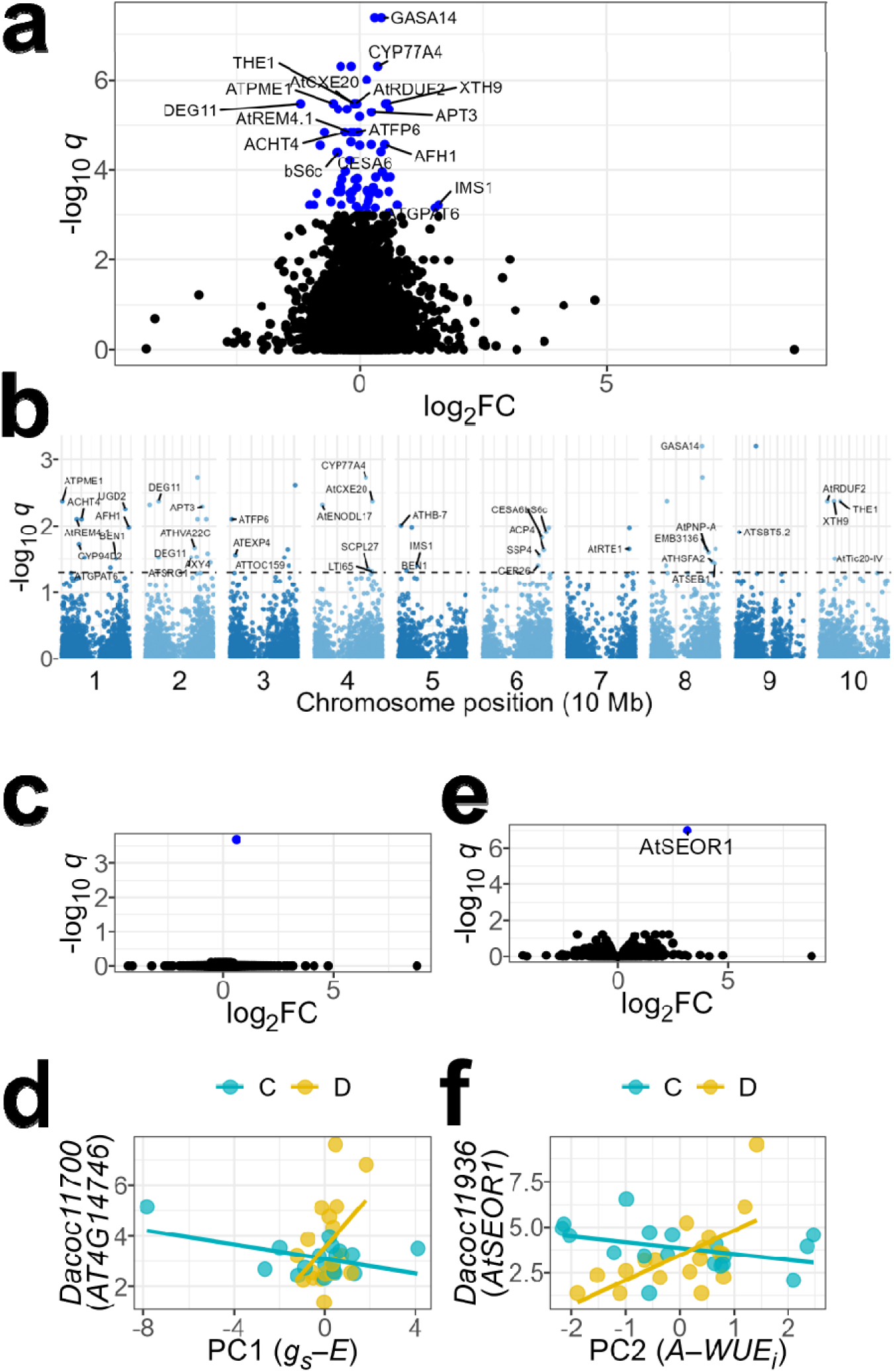
Volcano plots of –log_10_ *q*-values against log_2_ fold change for **(a)** the drought effect, **(c)** the isohydry effect (using PC1), and **(e)** the isohydry effect (using PC2) using likelihood ratio test (LRT) between a full model and a reduced model without the effect of concern. **(b)** Manhattan plot of –log_10_ *q*-values against chromosome position for the drought effect. **(d)** VST-transformed gene expression of Isohydry PC1-associated gene *Dacoc11700* (*AT4G14746*). **(f)** VST-transformed gene expression of Isohydry PC2-associated gene *Dacoc11936* (*AtSEOR1*) C and D denote well-watered control and drought treatment respectively.

For the *isohydry effect*, we only detected two genes that were significantly differentially expressed. One strongly associated with PC1 (*g_s_–E*) was Dacoc11700 (AT4G14746) (*q* = 0.025, **Figure 5c**), that encodes a neurogenic locus notch-like protein. Higher expression of Dacoc11700 led to a higher PC1 score, which implies steeper *g_s_–E* slopes and thus early stomatal closure, a traditional isohydric response (**Figure 5d**). Dacoc11700 was predicted to contain a signal peptide and locate on the outside of cell membrane. The gene strongly associated with PC2 (*A–WUE_i_*) was Dacoc11936 (SEOR1, Sieve Element Occlusion Related 1) (*q* = 0.00092, **Figure 5e**), that encodes a protein primarily located within the sieve elements in the phloem. Similarly, higher expression of SEOR1 led to a higher PC2 score (**Figure 5f**), which implies steeper *A–WUE_i_* slopes and thus decreased photosynthetic efficiency, also a traditional isohydric response.

### Differential transcript usage

We detected significant provenance-specific isoform switches between well-watered controls and drought-treated individuals for two provenances NP and THB, but not any overall effect of drought treatment or provenance alone.

For THB, we detected 2 isoform switches. Dacoc21458 (PRX52, Peroxidase 52) showed a significant isoform switch (*q* = 0.020) between a full functional isoform RB and an truncated isoform transcript7028 which had an alternative transcription start site with a missing signal peptide (**Figure 6a**). The isoform fraction of RB decreased from 98.89% to 41.60% between control and drought, whereas that of transcript7028 increased from 1.11% to 58.40%, constituting a change of 57.29% (**Figure 6b**). We also discovered a novel and previously unannotated gene 16703 on chromosome 7 that showed a significant isoform switch (*q* = 4.72e-15) (**Supplementary Figure 3a**), which had 4 non-coding isoforms of varied lengths and no domain or topology could be predicted. All 4 isoforms had relatively similar abundance in control, but the longest isoform transcript16686 dominated 94.30% of the abundance in drought, constituting a change in isoform fraction of 80.06% (**Supplementary Figure 3b**).

**Figure 6.**
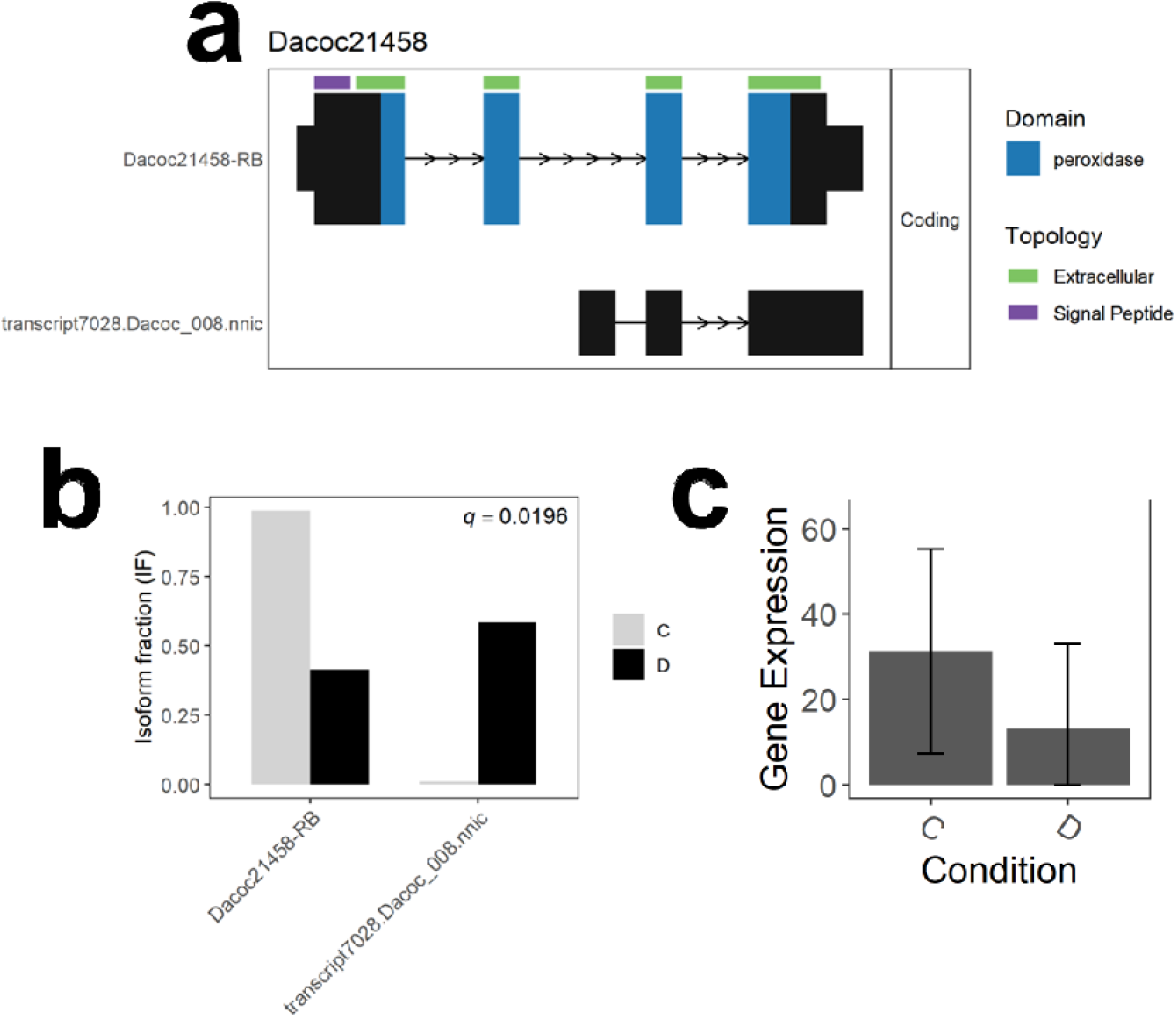
Isoform switch of Dacoc21458 (PRX52) between control and drought treatment in provenance THB. **(a)** Structures and annotations of the two isoforms. **(b)** Gene expression between control (C) and drought (D) conditions. **(c)** Isoform fraction of the two isoforms between control and drought conditions. **(c)** Gene expression between control (C) and drought (D) conditions.

For NP, we detected 2 further isoform switches. Dacoc26112 (ANN3, Annexin 3) showed a significant switch (*q* = 0.0056) between a full functional isoform RA and three other alternative isoforms (**Supplementary Figure 4a**). The isoform fraction of RA increased from 10.22% to 45.23% between control and drought, whereas the non-coding isoform transcript41999 decreased from 67.18% to 8.03% (**Supplementary Figure 4b**). Another gene Dacoc06197 (LTPG5, Non-specific lipid transfer protein GPI-anchored 5) switched between a full functional isoform RA and an alternative isoform transcript43524 with an earlier start site (*q* = 0.034) (**Supplementary Figure 5a**). The isoform fraction of RA increased from 55.62% to 94.59% between control and drought, whereas that of transcript43524 decreased from 44.38% to 5.41%, constituting a change of 38.98% (**Supplementary Figure 5b**).

However most importantly, these genes have no significant difference in gene-level expression between control and treatment, and would not have been detected in the differential gene expression analysis (**Figure 6c**, **Supplementary Figure 3c**, **Supplementary Figure 4c**, and **Supplementary Figure 5c**). Therefore, isoform switches are the sole mechanism that regulates the response to drought in these genes.

## Discussion

### Drought response and isohydry in Siamese rosewood

Decreased soil water content (SWC) imposes significant constraints on *g_s_* in Siamese rosewood, but the magnitude and direction of responses are provenance-dependent, as shown by the SWC × provenance interaction, which was significant for *g_s_* and *A*, and marginal for *E*. The near-orthogonality of *g_s_*–*E* and *A*–*WUE_i_*implies an independent carbon-economics dimension, in which adjustments in stomatal water-flux is decoupled from photosynthetic performance. This observation means that different provenances adopt distinct combinations of flux control and carbon maintenance in response to drought stress. Some provenances, such as THB exhibited sharp reductions in *E* with relatively modest declines in *A*. In contrast, individuals from KKH and DN decreased *E* in response to declining water availability but increased *A*, thus increasing *WUE_i_* even when the level of drought increases.

The *E*–*A* decoupling has been observed when a change in stomatal flux does not always correspond to a change in photosynthetic capacity. In some cases of extreme heat, transpiration continues even as photosynthesis declines to near zero (*E*↑*A*↓)^50^, which may stem from a passive physical effect of high temperatures that increases the fluidity of water and the permeability of guard cells^51^. This response also has an adaptive advantage where sacrificing water is necessary for cooling by transpiration to enhance the survival of leaves but ultimately will deplete water stores^52^. However, Siamese rosewood provenances KKH and DN display a rare, opposite relationship, where transpiration decreases and photosynthesis continues to increase (*E*↓*A*↑). There may be a completely different mechanism via mesophyll CO_2_ conductance, thus maintaining comparatively high CO_2_ supply to chloroplasts even as stomata are tightly closed^53^. Aquaporin-mediated CO_2_ diffusion has been found to modulate mesophyll conductance and to respond to stress and hormones, such as drought and ABA priming^54–56^. Mesophyll conductance is not measured in our study and its occurrence in Siamese rosewoods is speculative, however, the observation of elevated *WUE_i_* during drought for KKH and DN may be linked to improved sustainability and growth outcomes of trees^57^, especially for a species such as Siamese rosewood, which is fast growing, ecologically pioneering species with high water consumption^20^.

Pigment and growth responses indicate functional acclimation without large structural shifts over our experimental timeframe. We observed that Δ *Height* and *Chl_a_* exhibit treatment × provenance interactions, whereas *SLA*, *LDMC*, *TSS*, and *Starch* show no detectable treatment or interaction effects. Such patterns are broadly consistent with the notion that short-term drought primarily elicits adjustments in gas exchange, pigments, and hormonal signalling, while structural reconfiguration often requires longer or more severe stress and can vary with species and context^58^. The high-*WUE_i_*provenances, KKH and DN, continue to reflect the adaptive benefits of *E*–*A* decoupling, where in both provenances *Chl_a_* and Δ *Height* has increased in drought-treated samples.

Characterising isohydry in a *g_s_*–*E*/*A*–*WUE_i_*framework as developed in the present study depends on its definition. If isohydry is defined solely by the stomatal closure in declining water availability, then judging by the direction of *g_s_*–*E* in the provenance-PCA, PT and THA would be the most isohydric, and THB would be the most anisohydric at whole-provenance levels. If the definition of isohydry also considers the assimilation and water use efficiency, then judging by the direction of PC2 in the provenance-PCA, THA would be the most isohydric and KKH would be the most anisohydric at whole-provenance levels. Our study highlights the heterogeneity in drought response in line with contemporary views of the isohydry-anisohydry continuum, which emphasises that water-potential regulation emerges from multiple, only partly covarying stomatal, hydraulic, osmotic, and metabolic processes, rather than a single trait or score^59,60^.

### Wettest months, not dry, determine the isohydry-anisohydry continuum

Precipitation in wettest months is found to be the single determinant factor for the isohydry-anisohydry continuum in Siamese rosewood, with more isohydric the provenance associated with higher the precipitation. This may be explained by the high wet-season precipitation acting as a trait construction window which promotes anatomical features, such as larger, more conductive earlywood vessels^61^. However, larger vessels often track greater vulnerability to cavitation, that is, higher P50^62^, and thus requires stronger stomatal control to protect the vulnerable xylem^63^. Moreover, stomatal density and size and cuticle properties are gradually set towards leaf maturation^64^, which may peak after the wettest month. Higher precipitation is reported to increase stomatal density across plant communities^65^, and may also reduce cuticle thickness^66^. Thus, the development and growth in wettest months are likely to determine anatomical adaptation that predicts drought response.

An additional effect of higher precipitation in wettest months may be to reduce the carbon penalty commonly associated with isohydry during drought. Wet-season carbon gain can build up larger non-structural carbohydrate (NSC) pools, which are used for maintenance respiration, osmotic adjustment, and defence^67^. NSC might persist into the dry season such that carry-over carbon can buffer the risk of carbon starvation under stomatal closure^68^. A wetter peak month also boosts soil and plant water stores, and thus higher capacitance^69^. Stomata spend less total time closed to maintain the same hydraulic safety, thus the area under the photosynthetic curve can be larger even under isohydry^70,71^. In contrast, drought stresses in drier months only act on the anatomical and hydraulic settings after they are built in wetter months, and thus stomatal and hydraulic limits are unlikely to be constrained by the dry season.

### Differential gene expression reveals different pathways

Most of our understanding of water status regulation along an isohydry-anisohydry continuum in plants is at the level of hydraulics and stomatal physiology while very little is known of its genetic control. Comparative omics has begun to reveal drought-response modules in model and crop plants, and a few tree systems, but explicit searches for putative regulators of isohydry, beyond generic drought or ABA pathways, remain scarce and largely indirect^72,73^. Here we take a different tack by mapping transcript abundance onto composite axes of water-use and carbon strategies, thereby nominating candidate regulators of these key traits.

The Dacoc11936 gene, which is homologous to SEOR1, is the only gene found to be associated with the phenotypic variation along the *A–WUE_i_* axis. It encodes a structural phloem P-protein that polymerises and occludes sieve elements upon wounding^74^. Although live-imaging studies show that SEOR1 agglomerations do not significantly alter phloem flow under non-wounding conditions in *Arabidopsis thaliana*^75^, phloem protection could represent an adaptive response to recurrent mechanical stress in Siamese rosewood due to seasonally dry conditions of southeast Asian forests. This prediction is plausible given that assimilation can be feedback-limited by sink unloading and transport capacity as phloem unloading is largely convective with a diffusion component^76^. Therefore, it is plausible that SEOR1-medicated occlusion stabilises phloem transport, buffering carbon allocation and thus *A–WUE_i_*.

Dacoc11700 (AT4G14746) is the only gene found to be associated with the phenotypic variation along the *g_s_–E* axis. It is annotated as a “neurogenic locus notch-like protein” and remains poorly experimentally characterised in *Arabidopsis*, but it is predicted to be extracellular^77^. It contains a signal peptide and thus could hypothetically contribute to cell-cell or apoplastic signal perception of cues such as ABA. It represents a candidate regulator for isohydry that requires functional validation, such as a CRISPR/Cas9 knock-out or overexpression complemented by sub-cellular localisation analyses to verify its apoplastic targeting.

We have identified many drought-responsive genes that have been validated in previous research. For example, GASA14 is a small, cysteine-rich, secreted apoplastic peptide that integrates GA and ABA responses and modulates reactive oxygen species (ROS) accumulation^78^. DEG11 is a chloroplastic protease that helps degrade photodamaged photosystem during photoinhibition, which is important for limiting photooxidative damage with drought-induced stomatal closure and excess excitation^79^. CXE20 binds strigolactones (SLs) which impact root system architecture and mycorrhizal signalling, such that altered SL availability can shift water foraging and carbon allocation under drought stress^80^. PME1, XTH9, and THE1 all contribute to the modulation of cell wall organisation: PME1 is responsible for pectin de-esterification of homogalacturonan in the cell wall that changes pectin stiffness and porosity^81^; XTH9 remodels hemicellulose and drives xylem cell expansion^82^; THE1 responses to cell wall damage and transduces into hormonal response such as abscisic acid (ABA) production and wall remodelling^83^. The associated matrix polysaccharides are synthesised or methyl-esterified in the Golgi cis-cisterna before secretion and thus can adjust wall composition during drought^84^.

### Differential transcript usage is a potential mechanism of local adaptation

Alternative transcript usage can enable rapid loss- or gain-of-function under drought, by toggling coding potential, reshaping domains, exposing transcripts to nonsense-mediated decay, or adding or removing signal peptides that redirect secretion and subcellular localization, thereby tuning processes from guard-cell signalling to cell-wall and cuticle biogenesis^85–87^. With most plant drought transcriptomics aggregating data to the gene level, isoform diversity and its functional consequences are obscured^88^, and therefore population-specific or locally adapted mechanisms to distinct hydroclimates are missed.

We detected two isoform switches between control and drought treatment specific to the provenance THB, which was identified as the most anisohydric considering *gs–E* axis. We observed that PRX52 switched to a drought-dominant, loss-of-function isoform with a missing signal peptide. PRX52 is a class III apoplastic peroxidase involved in the synthesis of S units in interfascicular fibres during lignification^89^. Class III peroxidases characteristically possess an N-terminal signal peptide for entry into the secretory pathway^90^, and co-localise with laccases to lignifying wall domains during secondary cell-wall deposition^91^. Therefore, the drought-dominant isoform of PRX52 is likely to result in reduced lignin synthesis and higher cell extensibility, which might facilitate stomatal opening and mesophyll expansion at lower turgour^92^. Class III peroxidases also broadly modulate reactive oxygen species (ROS) homeostasis during stress responses^93^. Reduced secretion of PRX52 may dampen the apoplastic hydrogen peroxide (H_2_O_2_) burst, which is a proximate trigger of stomatal closure, and thus sustain a higher *g_s_*. Other class III peroxidases PRX4, PRX33, PRX34, and PRX71 have been shown in guard cells^94^. This discovery of reduced secretion in PRX52 regulating anisohydry opens novel avenues for drought research, which may be validated in future studies with ROS staining and imaging in guard cells.

We detected a gain-of-function phenomenon in the provenance NP, where both ANN3 and LTPG5 switched from non-coding or non-native isoforms, respectively, to the functional coding isoform during drought. Annexins are Ca²L-dependent phospholipid-binding proteins that relocalise to the plasma membrane when cytosolic Ca²L rises, participate in vesicle trafficking and exocytosis, and modulate ROS–Ca²L signalling in stress^95^. ANN3 specifically has been linked to eATP-regulated growth and vesicle polarity^96^. The drought-biased ATTS switch may thus upgreulate ANN3 protein to stabilise both the plasma membrane and exocytosis that maintains cell wall supply and repair^97^. On the other hand, GPI-anchored LTPGs localise to the outer leaflet of the plasma membrane and are required for exporting cuticular lipids and depositing suberin to the plant surface^98^. LTPG5 has confirmed roles in defence signalling, typical of many GPI-APs that interface with RLKs at the cell surface^99^. The drought-responsive LTPG5 full-length isoform may act by reinforcing the cuticle and reducing non-stomatal water loss.

### Implications for conservation and drought research

This study provides spatially explicit evidence for drought-responsive conservation of Siamese rosewood, which is crucial in times of intensifying drought and extreme weather conditions in their habitat. It enables climate-adjusted provenancing and assisted gene flow, such as matching seed sources to planting sites using a dual filter that combines each provenance’s position in the isohydry space with wettest-month precipitation of both source and target sites. High-*WUE*LJ, growth-maintaining provenances, such as KKH and DN, are prime candidates for dry, variable environments, but should be deployed in diverse, mosaic mixes to hedge risk and preserve genetic diversity. We present marker panels supported by understanding of underlying mechanism useful for drought tolerance screening and breeding, such as common variants responsible for cell wall, but other key drought-responsive genes and isoform switches also offer novel avenues in engineering drought-tolerant plants. Future drought research in Siamese rosewood and at large may include common garden validation of wettest-month precipitation in shaping isohydry behaviour. Our work could support several objectives: (1) extend measurements into mesophyll conductance and test aquaporin involvement in improving water-use efficiency; (2) integrate our genomic markers into genomic selection for water-efficient growth; and (3) run operational pilots that compare survival, growth, and hydrological impacts of provenance mixes under real reforestation settings. Siamese rosewood is a promising species that has pioneering ability suitable for forest landscape restoration. Conserving this critically endangered species will continue to realise its ecological and socioeconomic value and may set a valuable model for other threatened tropical tree species in Southeast Asia.

## Data availability statements

Sequence and meta data of the transcriptomic analyses have been deposited under NCBI BioProject PRJNA1378856, which contains all BioSample and Sequence Read Archive (SRA) accessions.

## Author contributions

T.H.H.: designed the study, conducted the drought experiment, processed the RNA samples, conducted the library preparation and sequencing, conducted the bioinformatic analyses, drafted the manuscript, and secured funding for the project;

K.L.: conducted the drought experiment and biochemical analyses and drafted the manuscript;

P.C.: collected the samples, revised the manuscript, and secured funding for the project;

V.C.: collected the samples, and revised the manuscript;

B.T.: collected the samples, revised the manuscript, and secured funding for the project;

R.J.: collected the samples, revised the manuscript, and secured funding for the project;

I.T.: collected the samples, revised the manuscript, and secured funding for the project;

J.J.M.: supervised the study, revised the manuscript, and secured funding for the project.

## Acknowledgements

This work is supported by funding to T.H.H., P.C., B.T., R.J., I.T., J.J.M. by the National Geographic Society (EC-95234R-22). We would like to thank Kate A. Hardwick from the Royal Botanic Gardens, Kew, for assistance in supplying samples from Thailand by V.C. via the Millenium Seed Bank Partnership. T.H.H. is supported with a Croucher Fellowship. T.H.H. wishes to personally thank the anonymous host of B612 for providing him a writing retreat and food during the winter of 2023 that initiates this manuscript.

## References

1. Martínez-Vilalta, J., Lloret, F. & Breshears, D. D. Drought-induced forest decline: causes, scope and implications. Biol Lett 8, 689–691 (2012).

2. Hammond, W. M. et al. Global field observations of tree die-off reveal hotter-drought fingerprint for Earth’s forests. Nat Commun 13, 1–11 (2022).

3. Gazol, A., Pizarro, M., Hammond, W. M., Allen, C. D. & Camarero, J. J. Droughts preceding tree mortality events have increased in duration and intensity, especially in dry biomes. Nature Communications 16, 1–11 (2025).

4. McDowell, N. G. et al. Mechanisms of woody-plant mortality under rising drought, CO2 and vapour pressure deficit. Nat Rev Earth Environ 3, 294–308 (2022).

5. Calvin, K. et al. IPCC, 2023: Climate Change 2023: Synthesis Report, Summary for Policymakers. Contribution of Working Groups I, II and III to the Sixth Assessment Report of the Intergovernmental Panel on Climate Change [Core Writing Team, H. Lee and J. Romero (Eds.)]. IPCC, Geneva, Switzerland. IPCC, 2023: Climate Change 2023: Synthesis Report. Contribution of Working Groups I, II and III to the Sixth Assessment Report of the Intergovernmental Panel on Climate Change [Core Writing Team, H. Lee and J. Romero (eds.)]. IPCC, Geneva, Switzerland. (2023). doi:10.59327/ipcc/ar6-9789291691647.001.

6. Alberto, F. J. et al. Potential for evolutionary responses to climate change – evidence from tree populations. Glob Chang Biol 19, 1645 (2013).

7. Brook, B. W., Sodhi, N. S. & Bradshaw, C. J. A. Synergies among extinction drivers under global change. Trends Ecol Evol 23, 453–460 (2008).

8. Mekong River Commission. Mekong Low Flow and Drought Conditions in 2019–2021: Hydrological Conditions in the Lower Mekong River Basin. https://www.mrcmekong.org/wp-content/uploads/2024/08/LowFlowReport20192021.pdf (2022).

9. Thirumalai, K., DInezio, P. N., Okumura, Y. & Deser, C. Extreme temperatures in Southeast Asia caused by El Ninõ and worsened by global warming. Nat Commun 8, 1–8 (2017).

10. Zhang, L., Chen, Z. & Zhou, T. Human Influence on the Increasing Drought Risk Over Southeast Asian Monsoon Region. Geophys Res Lett 48, e2021GL093777 (2021).

11. Hughes, A. C. Understanding the drivers of Southeast Asian biodiversity loss. Ecosphere 8, e01624 (2017).

12. Ang, W. J., Park, E., Pokhrel, Y., Tran, D. D. & Loc, H. H. Dams in the Mekong: a comprehensive database, spatiotemporal distribution, and hydropower potentials. Earth Syst Sci Data 16, 1209–1228 (2024).

13. UNODC. World Wildlife Crime Report: Trafficking in Protected Species. (United Nations Publication, 2016).

14. Barstow, M. et al. Dalbergia cochinchinensis. The IUCN Red List of Threatened Species 2022 https://www.iucnredlist.org/species/215342548/2822125 (2022).

15. Gaisberger, H. et al. Range-wide priority setting for the conservation and restoration of Asian rosewood species accounting for multiple threats and ecogeographic diversity. Biol Conserv 270, 109560 (2022).

16. CDRI. Community forestry for sustainable forest management and livelihhods: a case study of Osoam Community FOrest users group. Cambodia Development Review 18, (2014).

17. Luoma-aho, T., Hong, L. T., Ramanatha Rao, V. & Sim, H. C. Forest Genetic Resources Conservation and Management: Proceedings of the Asia Pacific Forest Genetic Resources Programme (APFORGEN) Inception Workshop. (2003).

18. APFORGEN. Conserving Rosewood Genetic Resources for Resilient Livelihoods in the Mekong - Project Inception Workshop Report. (2018).

19. Jalonen, R. et al. Effective Conservation of Asian Rosewoods (Dalbergias) and Their Genetic Resources Through Regional Collaboration. in Handbook of Asian Rosewoods (eds. Warrier, R. R., Joshi, G. & Arunkumar, A. N.) 461–471 (Springer, Singapore, 2025). doi:10.1007/978-981-96-9459-4_32/TABLES/1.

20. Hung, T. H. et al. Physiological responses of rosewoods Dalbergia cochinchinensis and D. oliveri under drought and heat stresses. Ecol Evol 10.1002/ece3.6744 (2020) doi:10.1002/ece3.6744.

21. McDowell, N. et al. Mechanisms of plant survival and mortality during drought: why do some plants survive while others succumb to drought? New Phytologist 178, 719–739 (2008).

22. Hartvig, I. et al. Population genetic structure of the endemic rosewoods *Dalbergia cochinchinensis* and *D. oliveri* at a regional scale reflects the Indochinese landscape and life-history traits. Ecol Evol 8, 530–545 (2018).

23. Hartvig, I. et al. Conservation genetics of the critically endangered Siamese rosewood (Dalbergia cochinchinensis): recommendations for management and sustainable use. Conservation Genetics 1–16 (2020) doi:10.1007/s10592-020-01279-1.

24. Hung, T. H. et al. Range-wide differential adaptation and genomic offset in critically endangered Asian rosewoods. Proc Natl Acad Sci U S A 120, e2301603120 (2023).

25. Gibson, G. The environmental contribution to gene expression profiles. Nature Reviews Genetics 2008 9:8 9, 575–581 (2008).

26. Ackermann, M., Sikora-Wohlfeld, W. & Beyer, A. Impact of Natural Genetic Variation on Gene Expression Dynamics. PLoS Genet 9, e1003514 (2013).

27. Benny, J., Pisciotta, A., Caruso, T. & Martinelli, F. Identification of key genes and its chromosome regions linked to drought responses in leaves across different crops through meta-analysis of RNA-Seq data. BMC Plant Biol 19, 1–18 (2019).

28. Benny, J. et al. Gaining Insight into Exclusive and Common Transcriptomic Features Linked to Drought and Salinity Responses across Fruit Tree Crops. Plants 9, 1–20 (2020).

29. Castillejo, M. A., Pascual, J., Jorrín-Novo, J. V. & Balbuena, T. S. Proteomics research in forest trees: A 2012-2022 update. Front Plant Sci 14, (2023).

30. Song, F. et al. Transcriptome and association mapping revealed functional genes respond to drought stress in Populus. Front Plant Sci 13, 829888 (2022).

31. Hung, T. H. et al. Reference transcriptomes and comparative analyses of six species in the threatened rosewood genus Dalbergia. Sci Rep 10, 17749 (2020).

32. Canham, C. D. & Murphy, L. The demography of tree species response to climate: seedling recruitment and survival. Ecosphere 7, e01424 (2016).

33. Hatfield, J. L. & Dold, C. Water-use efficiency: Advances and challenges in a changing climate. Front Plant Sci 10, 429990 (2019).

34. Pérez-Harguindeguy, N. et al. New handbook for standardised measurement of plant functional traits worldwide. Aust J Bot 61, 167–234 (2013).

35. Sims, D. A. & Gamon, J. A. Relationships between leaf pigment content and spectral reflectance across a wide range of species, leaf structures and developmental stages. Remote Sens Environ 81, 337–354 (2002).

36. Osaki, M., Shinano, T. & Tadano, T. Redistribution of carbon and nitrogen compounds from the shoot to the harvesting organs during maturation in field crops. Soil Sci Plant Nutr 37, 117–128 (1991).

37. Prjibelski, A. D. et al. Accurate isoform discovery with IsoQuant using long reads. Nature Biotechnology 2023 41:7 41, 915–918 (2023).

38. Love, M. I., Huber, W. & Anders, S. Moderated estimation of fold change and dispersion for RNA-seq data with DESeq2. Genome Biol 15, 1–21 (2014).

39. Ignatiadis, N., Klaus, B., Zaugg, J. B. & Huber, W. Data-driven hypothesis weighting increases detection power in genome-scale multiple testing. Nature Methods 2016 13:7 13, 577–580 (2016).

40. Wu, T. et al. clusterProfiler 4.0: A universal enrichment tool for interpreting omics data. The Innovation 2, 100141 (2021).

41. Korotkevich, G. et al. Fast gene set enrichment analysis. bioRxiv 060012 (2021) doi:10.1101/060012.

42. Vitting-Seerup, K., Sandelin, A. & Berger, B. IsoformSwitchAnalyzeR: analysis of changes in genome-wide patterns of alternative splicing and its functional consequences. Bioinformatics 35, 4469–4471 (2019).

43. Anders, S., Reyes, A. & Huber, W. Detecting differential usage of exons from RNA-seq data. Genome Res 22, 2008 (2012).

44. Kang, Y. J. et al. CPC2: a fast and accurate coding potential calculator based on sequence intrinsic features. Nucleic Acids Res 45, W12–W16 (2017).

45. El-Gebali, S. et al. The Pfam protein families database in 2019. Nucleic Acids Res 47, D427–D432 (2019).

46. Nielsen, H. Practical Applications of Language Models in Protein Sorting Prediction: SignalP 6.0, DeepLoc 2.1, and DeepLocPro 1.0. Methods in Molecular Biology 2941, 153–175 (2025).

47. Thumuluri, V., Almagro Armenteros, J. J., Johansen, A. R., Nielsen, H. & Winther, O. DeepLoc 2.0: multi-label subcellular localization prediction using protein language models. Nucleic Acids Res 50, W228–W234 (2022).

48. Hallgren, J. et al. DeepTMHMM predicts alpha and beta transmembrane proteins using deep neural networks. bioRxiv 2022.04.08.487609 (2022) doi:10.1101/2022.04.08.487609.

49. Erdos, G. & Dosztányi, Z. AIUPred: combining energy estimation with deep learning for the enhanced prediction of protein disorder. Nucleic Acids Res 52, W176–W181 (2024).

50. Urban, J., Ingwers, M. W., McGuire, M. A. & Teskey, R. O. Increase in leaf temperature opens stomata and decouples net photosynthesis from stomatal conductance in Pinus taeda and Populus deltoides x nigra. J Exp Bot 68, 1757–1767 (2017).

51. Marchin, R. M., Medlyn, B. E., Tjoelker, M. G. & Ellsworth, D. S. Decoupling between stomatal conductance and photosynthesis occurs under extreme heat in broadleaf tree species regardless of water access. Glob Chang Biol 29, 6319–6335 (2023).

52. Aparecido, L. M. T., Woo, S., Suazo, C., Hultine, K. R. & Blonder, B. High water use in desert plants exposed to extreme heat. Ecol Lett 23, 1189–1200 (2020).

53. Théroux-Rancourt, G., Éthier, G. & Pepin, S. Threshold response of mesophyll CO2 conductance to leaf hydraulics in highly transpiring hybrid poplar clones exposed to soil drying. J Exp Bot 65, 741–753 (2014).

54. Perez-Martin, A. et al. Regulation of photosynthesis and stomatal and mesophyll conductance under water stress and recovery in olive trees: correlation with gene expression of carbonic anhydrase and aquaporins. J Exp Bot 65, 3143–3156 (2014).

55. Flexas, J. et al. Tobacco aquaporin NtAQP1 is involved in mesophyll conductance to CO2in vivo. The Plant Journal 48, 427–439 (2006).

56. Chen, J. et al. Aquaporins and CO2 diffusion across biological membrane. Front Physiol 14, 1205290 (2023).

57. Rancourt, G. T., Éthier, G. & Pepin, S. Greater efficiency of water use in poplar clones having a delayed response of mesophyll conductance to drought. Tree Physiol 35, 172–184 (2015).

58. Hancock, R. D. et al. Physiological, biochemical and molecular responses of the potato (Solanum tuberosumL.) plant to moderately elevated temperature. Plant Cell Environ 37, 439–450 (2014).

59. Martínez-Vilalta, J. & Garcia-Forner, N. Water potential regulation, stomatal behaviour and hydraulic transport under drought: deconstructing the iso/anisohydric concept. Plant Cell Environ 40, 962–976 (2017).

60. Hochberg, U., Rockwell, F. E., Holbrook, N. M. & Cochard, H. Iso/Anisohydry: A Plant–Environment Interaction Rather Than a Simple Hydraulic Trait. Trends Plant Sci 23, 112–120 (2018).

61. Schreiber, S. G., Hacke, U. G. & Hamann, A. Variation of xylem vessel diameters across a climate gradient: insight from a reciprocal transplant experiment with a widespread boreal tree. Funct Ecol 29, 1392–1401 (2015).

62. Lobo, A. et al. Assessing inter- and intraspecific variability of xylem vulnerability to embolism in oaks. For Ecol Manage 424, 53 (2018).

63. Salvi, A. M. et al. Hydroscapes, hydroscape plasticity and relationships to functional traits and mesophyll photosynthetic sensitivity to leaf water potential in Eucalyptus species. Plant Cell Environ 45, 2573–2588 (2022).

64. Yang, K. tong, Chen, G. peng & Xian, J. ren. Stomatal distribution pattern for 90 species in Loess Plateau – Based on replicated spatial analysis. Ecol Indic 148, 110120 (2023).

65. Liu, C. et al. Relationships of stomatal morphology to the environment across plant communities. Nat Commun 14, 1–11 (2023).

66. Li, X. et al. A thinner jacket for frosty and windy climates? Global patterns in leaf cuticle thickness and its environmental associations. New Phytologist 248, 107–124 (2025).

67. Hartmann, H. & Trumbore, S. Understanding the roles of nonstructural carbohydrates in forest trees - from what we can measure to what we want to know. New Phytol 211, 386–403 (2016).

68. Scartazza, A., Moscatello, S., Matteucci, G., Battistelli, A. & Brugnoli, E. Seasonal and inter-annual dynamics of growth, non-structural carbohydrates and C stable isotopes in a Mediterranean beech forest. Tree Physiol 33, 730–742 (2013).

69. O’Grady, A. P., Eamus, D. & Hutley, L. B. Transpiration increases during the dry season: patterns of tree water use in eucalypt open-forests of northern Australia. Tree Physiol 19, 591–597 (1999).

70. Matheny, A. M. et al. Observations of stem water storage in trees of opposing hydraulic strategies. Ecosphere 6, 1–13 (2015).

71. Zhang, Y. J., Meinzer, F. C., Qi, J. H., Goldstein, G. & Cao, K. F. Midday stomatal conductance is more related to stem rather than leaf water status in subtropical deciduous and evergreen broadleaf trees. Plant Cell Environ 36, 149–158 (2013).

72. Song, F. et al. Transcriptome and association mapping revealed functional genes respond to drought stress in Populus. Front Plant Sci 13, 829888 (2022).

73. Dal Santo, S., et al. Distinct transcriptome responses to water limitation in isohydric and anisohydric grapevine cultivars. BMC Genomics 2016 17:1 17, 1–19 (2016).

74. Ernst, A. M. et al. Sieve element occlusion (SEO) genes encode structural phloem proteins involved in wound sealing of the phloem. Proceedings of the National Academy of Sciences 109, E1980–E1989 (2012).

75. Froelich, D. R. et al. Phloem ultrastructure and pressure flow: Sieve-Element-Occlusion-Related agglomerations do not affect translocation. Plant Cell 23, 4428–4445 (2011).

76. Ross-Elliott, T. J. et al. Phloem unloading in arabidopsis roots is convective and regulated by the phloempole pericycle. Elife 6, (2017).

77. Hooper, C. M., Castleden, I. R., Tanz, S. K., Aryamanesh, N. & Millar, A. H. SUBA4: The interactive data analysis centre for Arabidopsis subcellular protein locations. Nucleic Acids Res 45, D1064–D1074 (2017).

78. Sun, S. et al. GASA14 regulates leaf expansion and abiotic stress resistance by modulating reactive oxygen species accumulation. J Exp Bot 64, 1637–1647 (2013).

79. Kapri-Pardes, E., Naveh, L. & Adam, Z. The Thylakoid Lumen Protease Deg1 Is Involved in the Repair of Photosystem II from Photoinhibition in Arabidopsis. Plant Cell 19, 1039 (2007).

80. Roesler, K. et al. Arabidopsis Carboxylesterase 20 Binds Strigolactone and Increases Branches and Tillers When Ectopically Expressed in Arabidopsis and Maize. Front Plant Sci 12, 639401 (2021).

81. Sénéchal, F. et al. Tuning of Pectin Methylesterification: PECTIN METHYLESTERASE INHIBITOR 7 MODULATES THE PROCESSIVE ACTIVITY OF CO-EXPRESSED PECTIN METHYLESTERASE 3 IN A pH-DEPENDENT MANNER. Journal of Biological Chemistry 290, 23320–23335 (2015).

82. Kushwah, S. et al. Arabidopsis XTH4 and XTH9 Contribute to Wood Cell Expansion and Secondary Wall Formation. Plant Physiol 182, 1946–1965 (2020).

83. Bacete, L. et al. THESEUS1 modulates cell wall stiffness and abscisic acid production in Arabidopsis thaliana. Proc Natl Acad Sci U S A 119, e2119258119 (2022).

84. Harholt, J., Suttangkakul, A. & Scheller, H. V. Biosynthesis of Pectin. Plant Physiol 153, 384 (2010).

85. Tognacca, R. S. et al. Alternative splicing in plants: current knowledge and future directions for assessing the biological relevance of splice variants. J Exp Bot 74, 2251–2272 (2023).

86. Lam, P. Y., Wang, L., Lo, C. & Zhu, F. Y. Alternative Splicing and Its Roles in Plant Metabolism. International Journal of Molecular Sciences 2022, Vol. 23, Page 7355 23, 7355 (2022).

87. Laloum, T., Martín, G. & Duque, P. Alternative Splicing Control of Abiotic Stress Responses. Trends Plant Sci 23, 140–150 (2018).

88. Sarantopoulou, D. et al. Comparative evaluation of full-length isoform quantification from RNA-Seq. BMC Bioinformatics 2021 22:1 22, 1–24 (2021).

89. Fernández-Pérez, F., Pomar, F., Pedreño, M. A. & Novo-Uzal, E. The suppression of AtPrx52 affects fibers but not xylem lignification in Arabidopsis by altering the proportion of syringyl units. Physiol Plant 154, 395–406 (2015).

90. Lüthje, S. & Martinez-Cortes, T. Membrane-Bound Class III Peroxidases: Unexpected Enzymes with Exciting Functions. International Journal of Molecular Sciences 2018, Vol. 19, Page 2876 19, 2876 (2018).

91. Hoffmann, N., Benske, A., Betz, H., Schuetz, M. & Lacey Samuels, A. Laccases and Peroxidases Co-Localize in Lignified Secondary Cell Walls throughout Stem Development. Plant Physiol 184, 806–822 (2020).

92. Lima, T. R. A. et al. Lignin composition is related to xylem embolism resistance and leaf life span in trees in a tropical semiarid climate. New Phytologist 219, 1252–1262 (2018).

93. Li, S., Zheng, H., Sui, N. & Zhang, F. Class III peroxidase: An essential enzyme for enhancing plant physiological and developmental process by maintaining the ROS level: A review. Int J Biol Macromol 283, 137331 (2024).

94. Arnaud, D. et al. Cytokinin-Mediated Regulation of Reactive Oxygen Species Homeostasis Modulates Stomatal Immunity in Arabidopsis. Plant Cell 29, 543–559 (2017).

95. Davies, J. M. Annexin-Mediated Calcium Signalling in Plants. Plants 2014, Vol. 3, Pages 128-140 3, 128–140 (2014).

96. Xu, J. et al. ATANN3 Is Involved in Extracellular ATP-Regulated Auxin Distribution in Arabidopsis thaliana Seedlings. Plants 12, 330 (2023).

97. Koerdt, S. N., Ashraf, A. P. K. & Gerke, V. Annexins and plasma membrane repair. Curr Top Membr 84, 43–65 (2019).

98. DeBono, A. et al. Arabidopsis LTPG Is a Glycosylphosphatidylinositol-Anchored Lipid Transfer Protein Required for Export of Lipids to the Plant Surface. Plant Cell 21, 1230 (2009).

99. Ali, M. A., Abbas, A., Azeem, F., Shahzadi, M. & Bohlmann, H. The Arabidopsis GPI-Anchored LTPg5 Encoded by At3g22600 Has a Role in Resistance against a Diverse Range of Pathogens. Int J Mol Sci 21, 1774 (2020).

