## Supplementary Information for "Environmental and transcriptomic determinants of drought response in critically endangered Siamese rosewood"

Tin Hang Hung^1,2*^
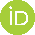
<https://orcid.org/0000-0001-9853-2053>, Kalyani Lenton^1^, Phourin Chhang^3^, Voradol Chamchumroon^4^
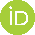
<https://orcid.org/0009-0004-0653-6137>, Bansa Thammavong^5^, Riina Jalonen^6^
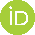
<https://orcid.org/0000-0003-1669-9138>, Ida Theilade^7^
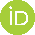
<https://orcid.org/0000-0003-3502-1277>, John J. MacKay^1,*^
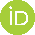
<https://orcid.org/0000-0002-4883-195X>

1. Department of Biology, University of Oxford, Oxford OX1 3EL, United Kingdom
2. Museum of Climate Change, The Chinese University of Hong Kong, Hong Kong
3. Institute of Forest and Wildlife Research and Development, Phnom Penh, Cambodia
4. The Forest Herbarium, Department of National Park, Wildlife and Plant Conservation, Ministry of Natural Resources and Environment, Bangkok 10900, Thailand
5. National Agriculture and Forestry Research Institute, Forestry Research Center, Vientiane, Laos
6. Bioversity International, 43400 UPM Serdang, Malaysia
7. Department of Geosciences and Natural Resource Management, University of Copenhagen, Rolighedsvej 23, 1958 Frederiksberg C, Denmark

* Corresponding authors:

Tin Hang Hung

John J. MacKay **Supplementary Table 1.** Details of the 210 trees sampled in this study.

*See separate spreadsheet*

**Supplementary Table 2.** List of all traits included in this study.

*See separate spreadsheet*

**Supplementary Table 3.** ANOVA tables of all traits with the effects of soil water content (*SWC*), provenance, and their interaction.

*See separate spreadsheet*

**Supplementary Table 4.** Differentially expressed genes on the effect of drought, only showing the significant ones (*P* < 0.05, Benjamini-Hochberg correction)

*See separate spreadsheet*

**Supplementary Table 5.** Gene ontology enrichment (GO) of the differentially expressed genes on the effect of drought, only showing the significant ones (*Q* < 0.05, Benjamini-Hochberg correction)

*See separate spreadsheet*

**Supplementary Figure 1. (a)** Change in leaf number (*Δ Leaf*), **(b)** change in branch number (*Δ Branch*), **(c)** anthocyanin content (*Ac*), **(d)** chlorophyll b content (*Chl_b_*), **(e)** carotenoid content (*Carot*), **(f)** leaf dry matter content (*LDMC*), **(g)** specific leaf area (*SLA*), **(h)** total soluble sugar content (*TSS*), and **(i)** starch content (*Starch*) between control (C) and drought (D) among six provenances.

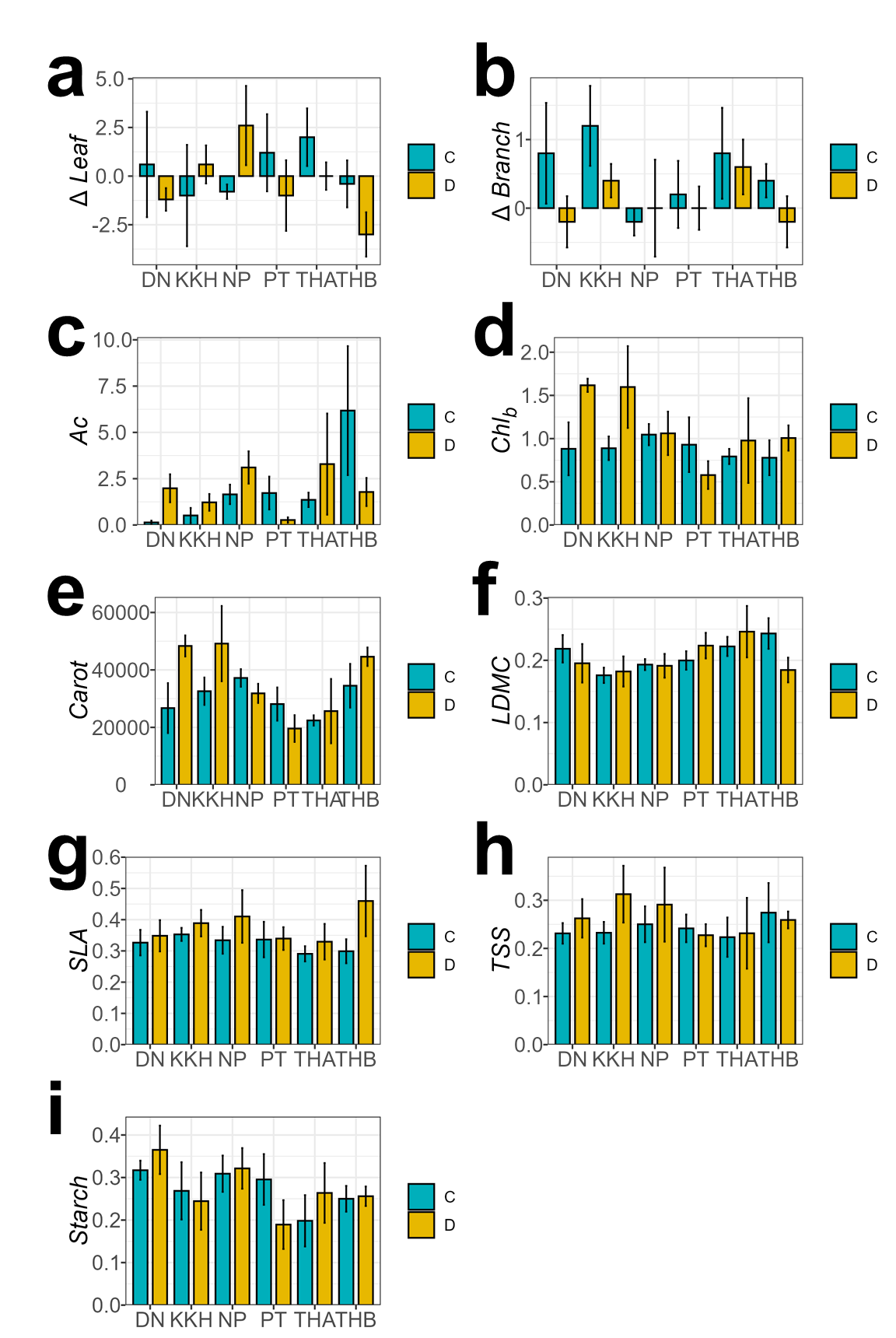

**Supplementary Figure 2.** Isoform switch of a novel gene Dacoc_007_16703 between control and drought treatment in provenance THB. **(a)** Structures and annotations of the four non-coding isoforms. **(b)** Isoform fraction of the two isoforms between control and drought conditions. **(c)** Gene expression between control (C) and drought (D) conditions.

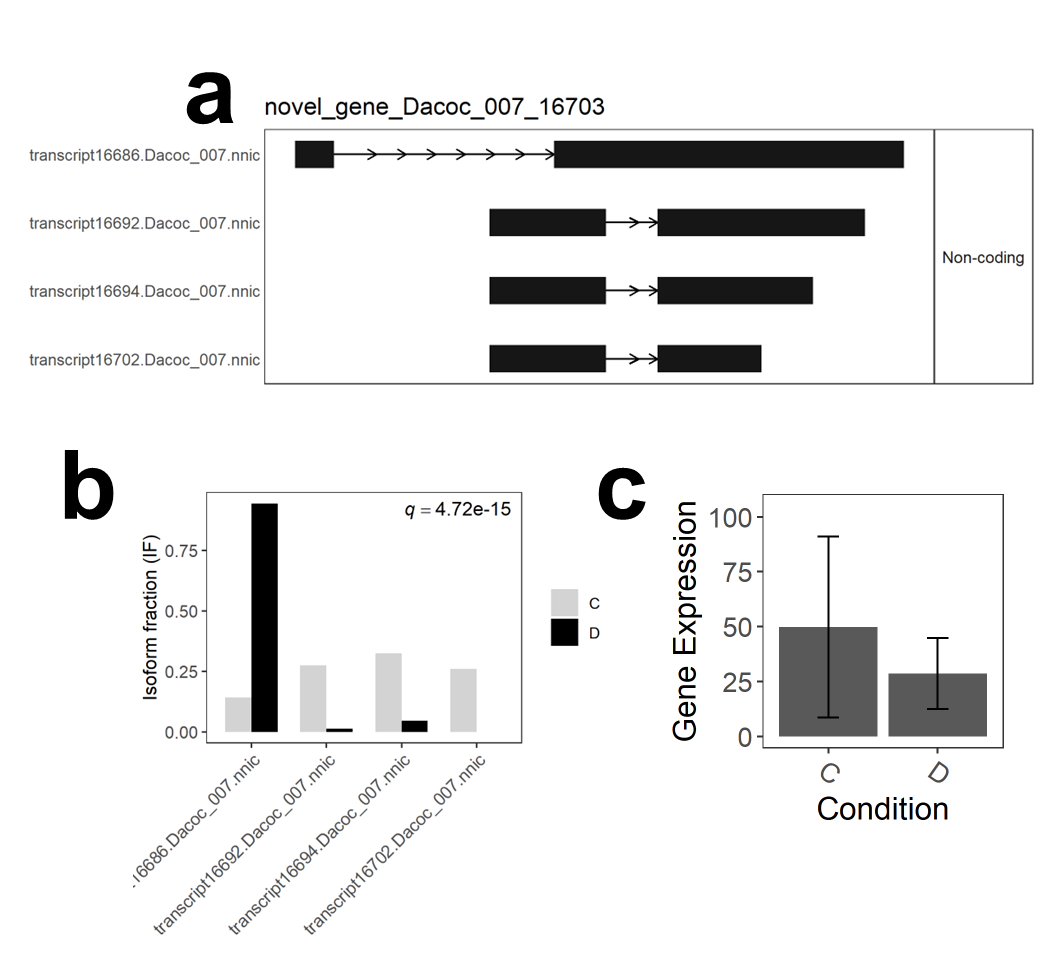

**Supplementary Figure 3.** Isoform switch of Dacoc26112 (ANN3) between control and drought treatment in provenance NP. **(a)** Structures and annotations of the three coding and one non-coding isoforms. **(b)** Isoform fraction of the two isoforms between control and drought conditions. **(c)** Gene expression between control (C) and drought (D) conditions

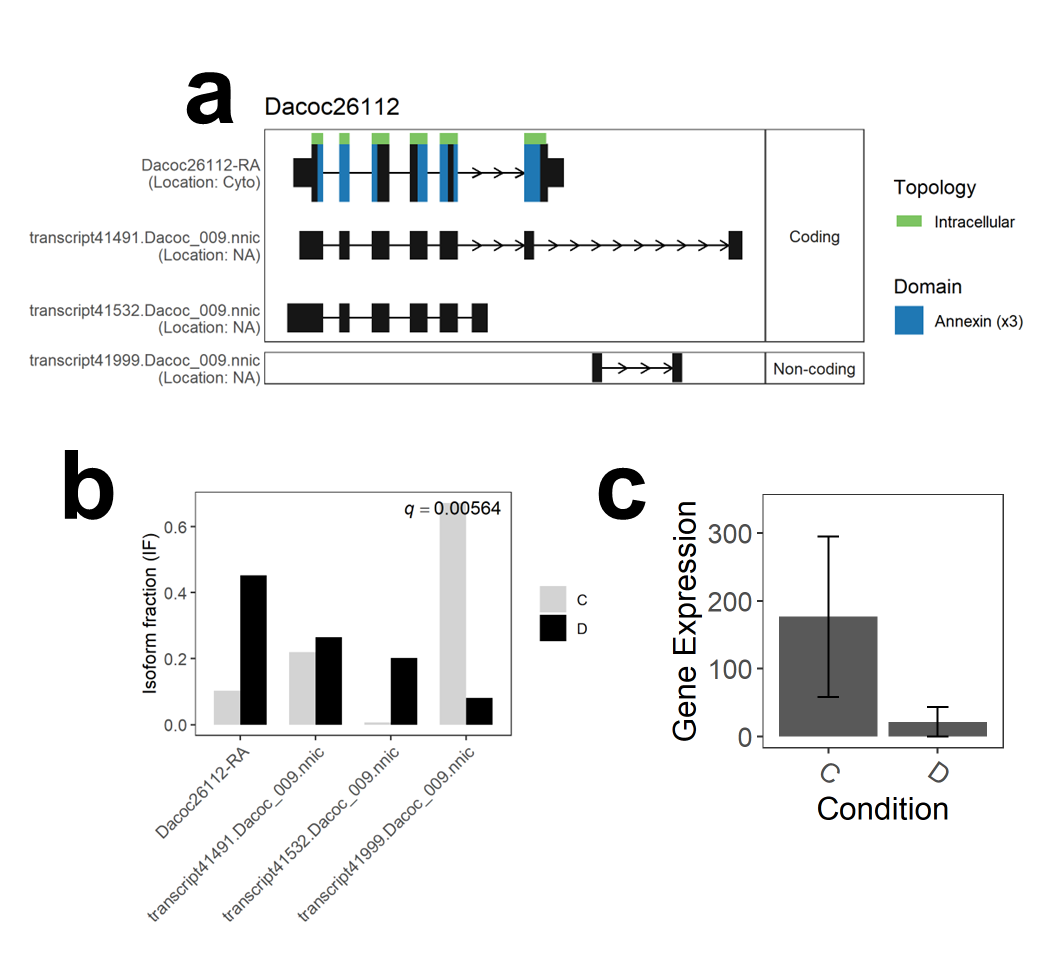

**Supplementary Figure 4.** Isoform switch of Dacoc06197 (LTPG5) between control and drought treatment in provenance NP. **(a)** Structures and annotations of the two coding isoforms. **(b)** Isoform fraction of the two isoforms between control and drought conditions.

**(c)** Gene expression between control (C) and drought (D) conditions.

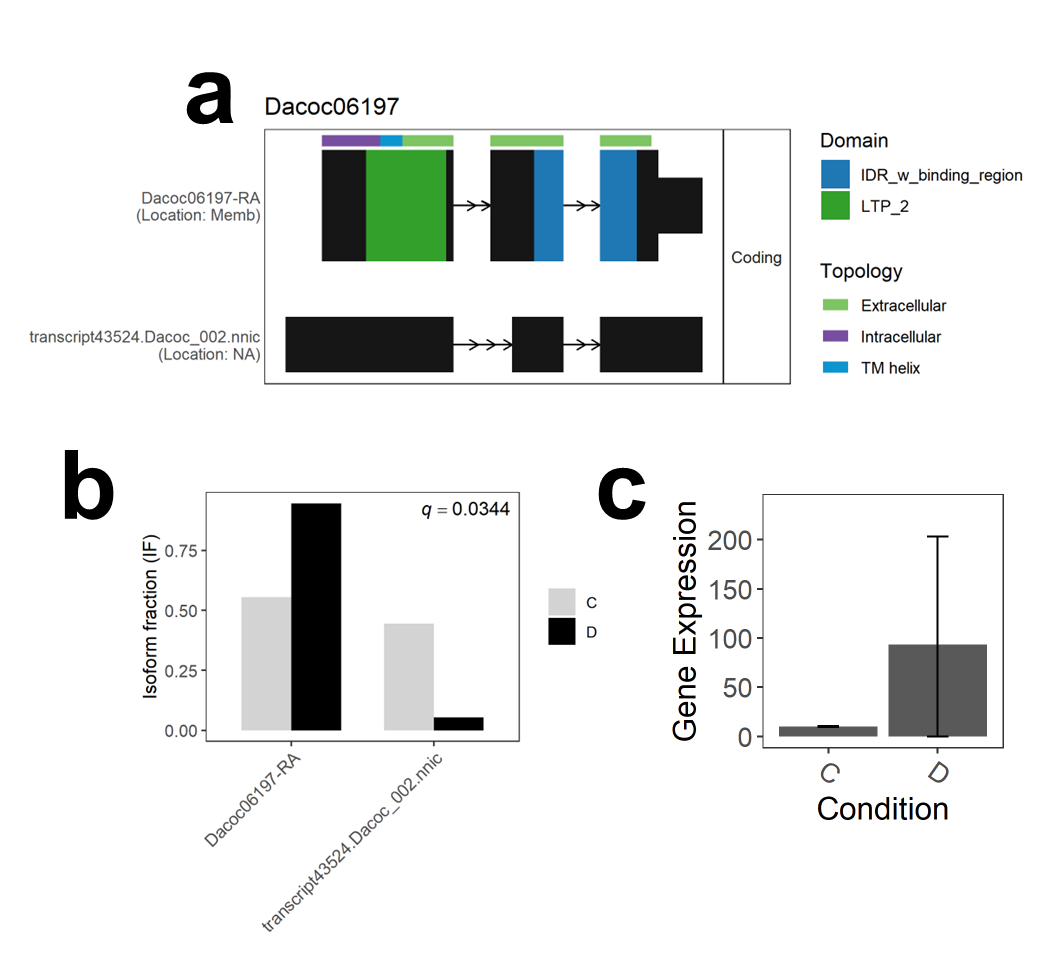
